# Facial gestures are enacted via a cortical hierarchy of dynamic and stable codes

**DOI:** 10.1101/2025.03.03.641159

**Authors:** Geena R. Ianni, Yuriria Vázquez, Adam G. Rouse, Marc H. Schieber, Yifat Prut, Winrich A. Freiwald

## Abstract

Successful communication requires the generation and perception of a shared set of signals. Facial gestures are one fundamental set of communicative behaviors in primates, generated through the dynamic arrangement of dozens of fine muscles. While much progress has been made uncovering the neural mechanisms of face perception, little is known about those controlling facial gesture production. Commensurate with the importance of facial gestures in daily social life, anatomical work has shown that facial muscles are under direct control from multiple cortical regions, including primary and premotor in lateral frontal cortex, and cingulate in medial frontal cortex. Furthermore, neuropsychological evidence from focal lesion patients has suggested that lateral cortex controls voluntary movements, and medial emotional expressions. Here we show that lateral and medial cortical face motor regions encode both types of gestures. They do so through unique temporal activity patterns, distinguishable well-prior to movement onset. During gesture production, cortical regions encoded facial kinematics in a context-dependent manner. Our results show how cortical regions projecting in parallel downstream, but each situated at a different level of a posterior-anterior hierarchy form a continuum of gesture coding from dynamic to temporally stable, in order to produce context-related, coherent motor outputs during social communication.

## Introduction

Faces are central to primate lives. Their visual appearance reveals a wide range of information to others; some through their structural properties, like age or identity, and others through their dynamic properties, like movement. Facial movements constitute a fundamental class of social communication signals in primates^1^. Nonverbal gestures can instantly convey hidden affective states^2^, the valence of external stimuli, as well as complex concepts like intention, social rank, and receptivity to future interactions^3^. Recent years have seen advances in the understanding of the neural mechanisms that analyze visual inputs to extract face-related information: a network of areas has been identified, each specialized in the processing of faces and each generating a distinct code for analyzing facial structure^4^. Recently, an area has been found containing cells specialized for the fine-grain analysis of facial dynamics^5–7^.

In human and non-human primates, the network for face perception is complemented by a motor network for the generation of facial gestures. These networks enable social dialogue. However, little is known about the neural mechanisms underlying facial movements in general, and facial gestures in particular. Past anatomical work has identified multiple cortical regions, including primary motor, ventral premotor, cingulate, and supplementary motor cortices, projecting directly to the facial nucleus in the pons^8–10^, which houses the motoneurons controlling facial muscles. This pattern implies that similar to the visual analysis of faces, the control of facial movements involves multiple cortical areas, and that similar to the control of dexterous behavior which has evolved specialized direct cortico-spinal projections^11^, cortical involvement in facial movement control is intimate.

Here we conduct the first neurophysiologic study to determine and compare the functional properties of cortical areas controlling facial movement. We leveraged two approaches: functional localization with magnetic resonance imaging (fMRI) and simultaneous multi-area, multi-channel recordings. By combining these techniques in an old-world monkey model system, which shares the highly differentiated facial musculature characteristic of humans^12–14^, we extracted single cell properties and population codes for facial gesture production during naturalistic social communication.

We show that in contrast to the neuropsychological hypothesis of medial-lateral functional segregation into emotional and voluntary control systems, all cortical face motor regions encode both types of gestures. Instead, the functional dissociation in the facial motor system is enacted by regionally-distinct temporal mechanisms, forming a continuum of stability in the code for facial gestures across cortex. Naturalistic facial communication is represented throughout the cortical face motor network, where hierarchy is in the temporal domain.

### Naturalistic social communication in the laboratory elicits distinct facial gestures

Much like humans, macaque monkeys use facial gestures to navigate the strict social hierarchies that define their large troops, both in captivity and in the wild^15–17^, providing an opportunity to investigate facial gesture production in an ecologically-valid model. In order to serve their communicative purpose, facial gestures must be distinct such that a receiver can recognize each and select a response. We focused on three gestures: lipsmack, threat, and chew. *Lipsmack*, an affiliative expression that involves rapid lip puckering accompanied by a flattening of the ears against the skull, can communicate receptivity to cooperative interactions as well as submission between conspecifics of unequal rank^18,19^. *Threat* is a communicative expression of adversarial nature: while staring straight ahead the jaw opens to display the canines. Its production may signal upcoming competitive interactions^20,21^. *Chew*, though not a social gesture, is a stereotyped, volitional ingestive movement. These movements require coordinated muscle activities generating complex visual patterns, which then are processed by the receiver^13,22,23^.

To elicit communications signals reliably, we developed a naturalistic social interaction paradigm. Ethologically-meaningful gestures were elicited via dynamic visual stimuli (e.g., movies of conspecifics, an interactive avatar, real-life interactions with conspecifics or experimenters, see Methods). Blocks of these “visuo-social” settings were interleaved pseudo randomly with blocks in which ingestive movements were elicited with food, and rest blocks in which the subject sat quietly. High-resolution video captured ongoing behavior synchronized to neural activity and stimulus display (**Figure 1A**). Offline, we used markerless tracking^24^ of facial landmarks in combination with manual scoring to determine movement onset, gesture type, and subsequently aligned all neural activity, recorded simultaneously, to facial gesture onsets (see *Methods*).

**Figure 1:**
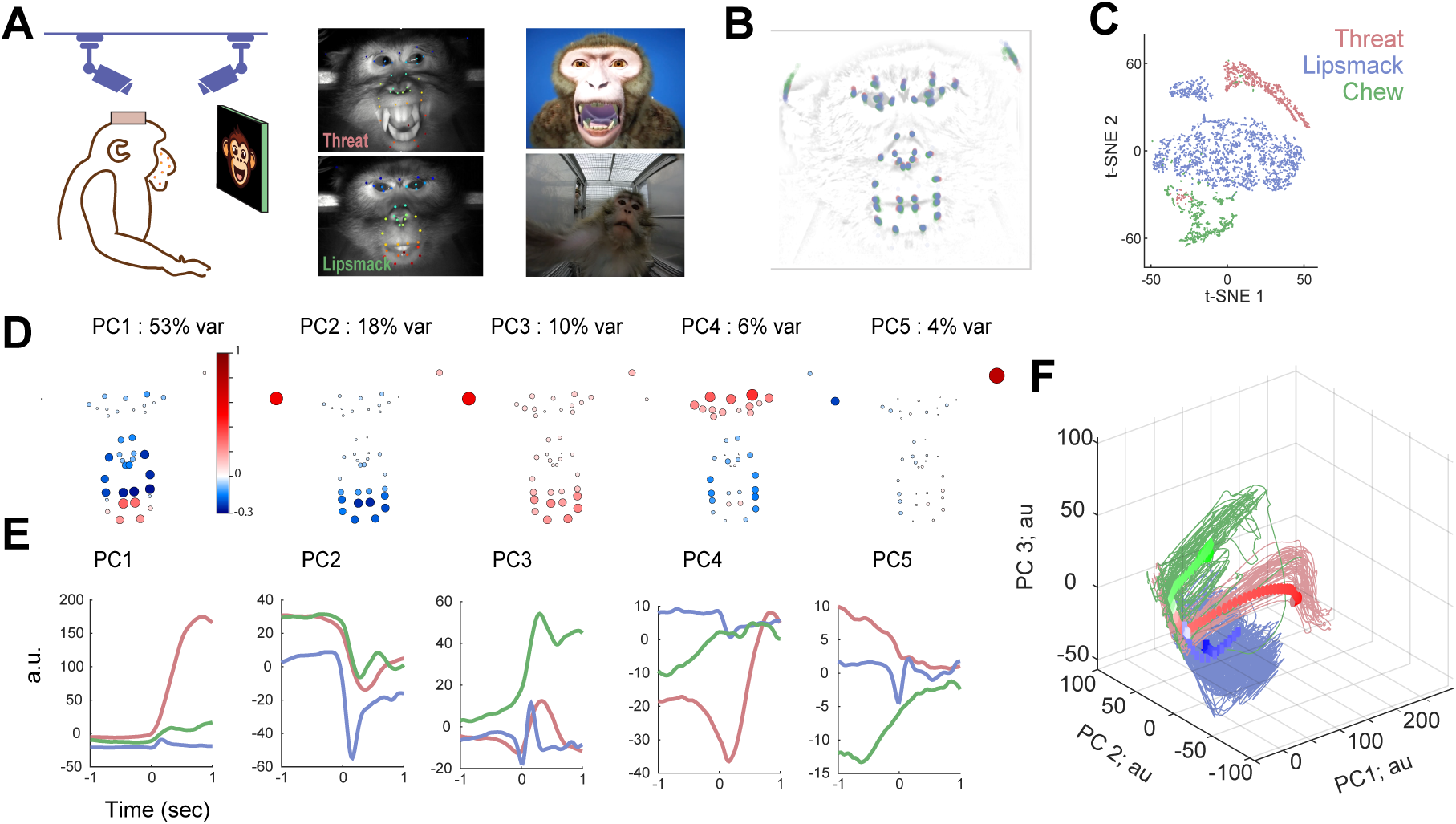
Facial gestures are distinguishable on a per-trial basis during a naturalistic social paradigm. A. Behavioral Paradigm. Subjects produced ethological facial gestures including affiliative lipsmack (blue), adversarial threat (red), and ingestive, non-social chew (green). High-resolution video captured ongoing behavior; example peak frames are shown (middle); overlay depicts locations of facial markers tracked continuously in two-dimensions using DeepLabCut. Socially communicative gestures were elicited with dynamic visuosocial stimuli such as an interactive monkey avatar (right, top), movies of conspecifics (right, bottom), and real-life interactions. All behavioral, neural, and stimulus data were synced to a common neural clock via TDT RZ2 BioAmp Processor. B. Initial facial posture at 1000 ms prior to movement onset. Each dot indicates 1 trial, 20% of trials are plotted, over an opacified depiction of subject’s face. C. Behavioral clustering by gesture-type. Considering gestures preceded by an adequate period of stillness, we extracted the two-dimensional positions of forty (subject 1) or fifty (subject 2) continuously-tracked facial markers (DeepLabCut) in the perimovement period (−500 to +100 milliseconds around movement onset). Data was binned at 100 ms, smoothed with a 50 ms kernel, and PCA applied to extract the top *N* components accounting for 95% of behavioral variance. These components were then reduced to two dimensions using tSNE and plotted; each dot represents an individual timepoint during an occurrence of threat (red), lipsmack (blue), or chew (green). One typical day of behavior is shown. D, F: Facial gestures trace stereotypical trajectories in behavioral space. We extracted the two-dimensional positions of markers over a longer period (−1000 to +1000 milliseconds around movement onset, re-binned at 10 ms, smoothed with 50 ms kernel). The top five facial gesture principle components (D, E) routinely accounted for >90% of behavioral variance, and varied systematically across gesture type. Marker weights depicted in original 2D face space (D) show gestures were composed of both shared and distinct movement components. Facial gesture movement patterns were stereotyped and readily distinguishable from one another on a per-trial basis (F); thin lines indicate individual trials, heavy lines indicate gesture-specific trial averages. One typical day of behavior is shown.

To serve as communicative signals, facial gestures should form distinct visual classes. We considered threat, lipsmack, and chew events preceded by an adequate period of stillness (see *Methods*, **Figure 1B**). We extracted the two-dimensional positions of facial markers throughout the movement period, and applied dimensionality reduction (PCA) to extract the top *N* components accounting for 95% of variance. Projecting the data on the first two components (**Figure 1C**) revealed distinct clusters for each gesture for both subjects, confirming that without formal training, our naturalistic paradigm produced discrete behaviors.

We found that the extracted components were related to movement in partially overlapping but broad regions of the face. Specifically, more than half the variance was explained by the first principal component (**Figure 1D**), mostly capturing movements of the upper and lower mouth. Principal components two and three captured additional modes of the lower face and left ear, the fourth PC captured movements of eyes and eye brows. All of these components varied systematically across gesture type (**Figure 1E**) such that distinct gestures were composed of shared and distinct effector activities (i.e., coordinated movement of a specific part of the face). Finally, we examined the temporal evolution of facial movement components (**Figure 1F** gesture averages in thick, individual trials in thin) and found unique gesture patterns that were stereotyped and readily distinguishable from one another on a per-trial basis.

### Simultaneous recording in functionally-localized medial and lateral face motor cortices during facial gesture production

While facial movements appear stereotypical and near reflexive (motor attributes which are often interpreted as signs for subcortical pattern generators, as is the case for respiration^25^ and mastication^26^), in humans cortical integrity is a prerequisite for these signals’ production^27^. Neuroanatomical studies have identified several cortical regions including primary motor, premotor, cingulate, and supplementary motor cortices containing direct projections to the facial nucleus in the pons^8–10^, which houses motoneurons controlling facial muscles (**Figure 2A)**.

**Figure 2:**
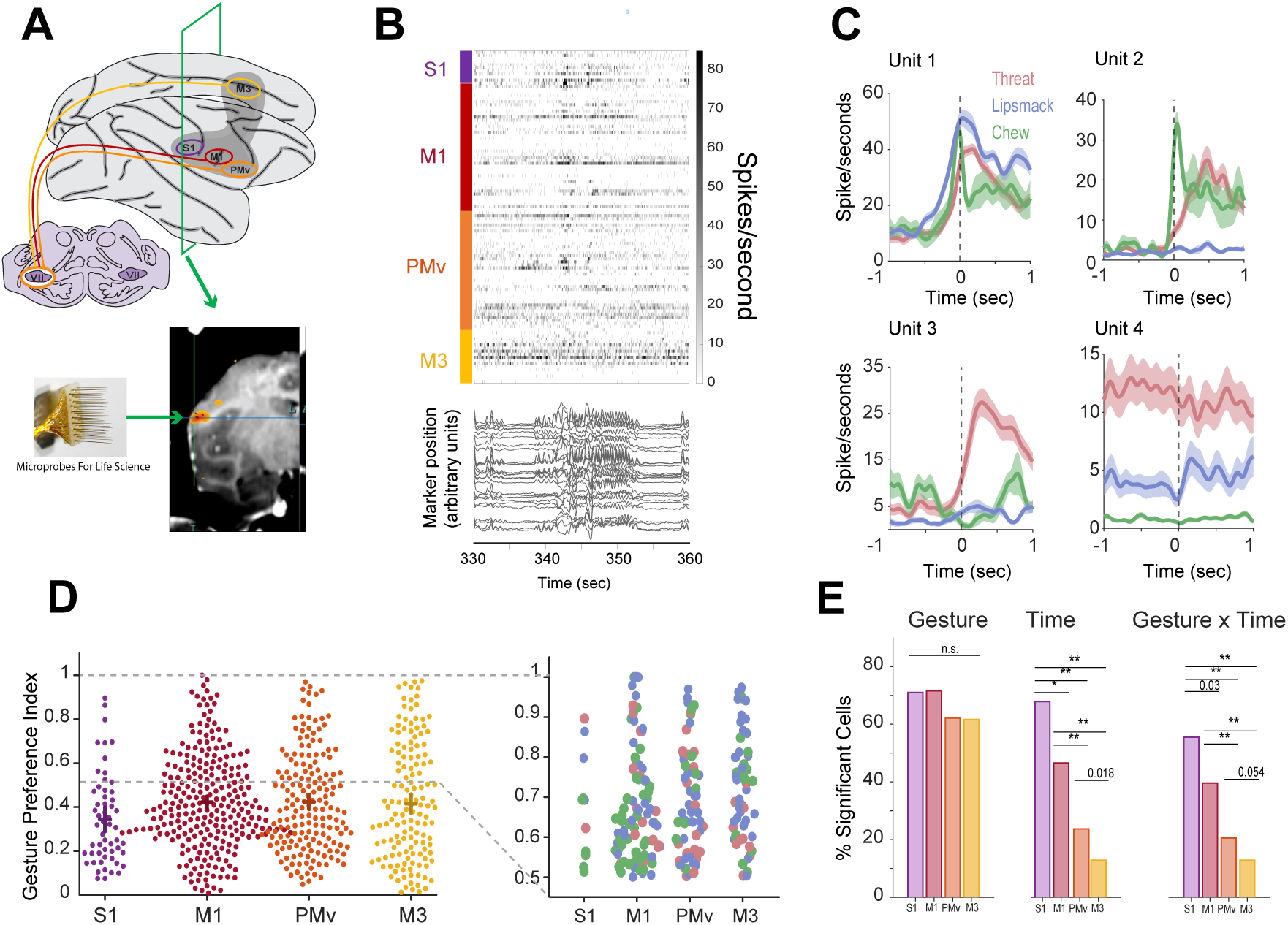
Electrophysiologic Activation of Single Cells in Cortical Face Motor Regions during Facial Gestures. A. FMRI-targeting for floating microelectrode array placement. We used an integrated functional/structural stereotaxic planning and intra-surgical tracking system (cortEXplore GmbH, Linz Austria^R^) to target four functionally-localized face motor areas with floating microelectrode arrays (FMA) for chronic electrophysiologic recordings. Targeted recording locations are depicted schematically (left) along with their known anatomical connectivity to facial nucleus; on the lateral surface: primary somatosensory cortex (S1; in purple), primary motor cortex (M1; red), ventral premotor cortex (PMv; orange) and on the deep medial surface, cingulate motor cortex (M3, yellow; note). Note all targeted regions were in the right hemisphere for all subjects. Each array consisted of 16-32 channels, yielding 32 channels in S1, 64-96 in M1, 32-64 in PMv, and 48-64 in M3, per subject. Electrode lengths were customized to maximize gray matter neural activity recordings. Insert depicts functional localization of M1 used for targeting. B. Raster of all simultaneously recorded single cells and behavior in the cortical face production network during ∼30 seconds of naturalistic facial movement. Cells are grouped by cortical region (top to bottom: S1, M1, PMv, M3). Below, tracings represent facial marker positions over time as subject engages in social interaction and produces facial gestures. Spike counts and behavior are binned at 50 ms; no additional alignment between neural activity and behavior was performed. C. Four example peri-event firing rates from the cortical face motor system. Spikes were aligned hand-scored gesture onsets, binned at 1 ms, and smoothed with 50 ms Gaussian kernel. Heavy trace indicates mean, shaded bars are 2 x SEM of trial-averaged response. Cells were chosen to highlight the diversity of modulations during production of three prototypical facial gestures. Cells in all implanted regions showed various degrees of tuning to facial gesture type; response shapes and latencies were often heterogeneous even within a single cell. D. Gesture preference indices (GPI), for all cells within each region (left). Hash marks represent region means, and their respective confidence intervals computed at 97.5% confidence level. A preference index of 1 indicates firing for one gesture exclusively, while and index of 0 indicates equal firing for all three gestures tested. A one-way ANOVA to compare the effect of cortical region on cells’ FEPI was non-significant (F(3,638) = [1.9515], p = 0.1201). Insert at right, distribution of GPIs for all selective cells defined as GPI > 0.5 per region, colored by preferred gesture type (lipsmack in blue, threat in red, chew in green). No region contained an overrepresentation of cells selective for one particular gesture. E. Fractions of cells in each region which modulated their firing rates on the basis of gesture type, time, or the interaction of gesture type with time. Spike counts on each trial were aligned to hand-scored movement onset, and binned in the 500 milliseconds prior to onset, and 500 milliseconds after onset. Spike counts were entered into a 2-way ANOVA (gesture type; 3 levels: threat, lipsmack, chew; time; 2 levels: before, after onset) and significance threshold set at p < 0.05 (multiple comparison corrected with Tukey’s HSD test). Large fractions of cells modulated by behavior type in all regions (S1: 71%, M1: 72%, PMv: 62%, M3: 62%). Chi-Square test for equality proportions revealed no effect of region on the fraction of cells modulated by behavior type; all regions contained similar fractions (*X*2 (3, *N* = 652) = 6.6449, *p* = 0.0841). There was a significant main effect of region on the fraction of cells modulated by time (S1: 68%, M1: 47%, PMv: 24%, M3: 14%; *X*2 (3, *N* = 652) = 84.0113, *p*=4.23×10^−18^); post-hoc comparisons (Bonferroni corrected) revealed significant differences in the fractions of time-modulated cells for all inter-areal comparisons. We found the same for the fraction of cells modulated by the interaction of time x gesture-type; post-hoc comparisons were significant for all inter-areal comparisons.

To dissect the contributions of these regions to facial gesture production, we developed an integrated functional/structural MRI-based stereotaxic planning and intra-surgical tracking approach (now cortEXplore GmbH, Linz Austria^R^, see *Methods*) to target four fMRI-localized regions in this network with chronic microelectrode arrays (MicroProbes for Life Sciences); areas included, on the lateral surface, primary motor (M1; red, maroon, pink), and ventral premotor cortex (PMv; blue, orange) and on the deep medial surface, cingulate motor cortex (M3, green and yellow). Additionally, we recorded from primary somatosensory cortex (S1; purple) due to its role in orofacial behaviors^28–31^. In what follows, we will refer to all of these regions as face motor regions. Each array consisted of 16 or 32 channels, resulting in per-subject totals of 64-96 channels in M1, 32-64 in PMv, 48-64 in M3, and 32 in S1, respectively (**Figure 2A**; see *Methods*). All arrays yielded numerous well-isolated single units over multiple recording days.

### Face motor regions contain similar mixtures of gesture-specific and gesture-general neurons

To test whether cortical regions are engaged in naturalistic facial movements, we examined movement-related activity of neurons and found cases of robust activity locked to facial movements evident in raw data traces (**Figure 2B)**. Neurons in all cortical face motor regions modulated their firing rates during production of all three facial gestures, often in a gesture-specific manner. **Figure 2C** shows the peri-event time histograms of four sample neurons with different response properties. Unit 3 increased its firing rate five-fold relative to a no-movement baseline exclusively during threats. Other neurons such as Unit 2, sharply increased activity for some gestures (chew), and more slowly others (threat) or not at all (lipsmacking). Unit 1’s activity increased prior to movement onset of all three gestures, with gesture-specific activity during the early movement phase. Finally, neurons such as Unit 4 showed marked activity differences for the three movement types without significant modulation locked to gesture onset.

To capture the relationship of neural activity to movement, we first quantified the fraction of cells significantly modulated by facial gesture production (see *Methods*). Large fractions of cells modulated by behavior type in all regions (**Figure 2E**, left distribution; S1: 71%, M1: 72%, PMv: 62%, M3: 62%; Chi-square test for equality of proportions: *χ*^2^(3, N = 652) = 6.6449, p = 0.084). Though gestures may appear reflexive, their production robustly engages neurons of multiple cortical regions.

A well-accepted neuropsychological schema of facial movement control is the dissociation of socio-emotional and volitional facial movements enacted via medial and lateral cortical motor representations, respectively^32–39^. This framework makes a strong prediction that the medial region (M3) contains an overrepresentation of cells strongly modulated by a specific type of facial gesture, while the lateral regions (M1, PMv) contain cells modulated by another.

To test this hypothesis, we defined for each cell the strength of neural tuning to gesture type and tested for differences between cortical areas. Tuning strength was quantified by gesture preference index, which is 0 if the cell fires equally for all gestures, and 1 if a cell fires for one gesture exclusively^40^. A one-way ANOVA to examine the effect of cortical region on the gesture selectivity index was non-significant (F3,638) = [1.9515], p = 0.1201): both medial and lateral regions contained a mixture of both highly gesture-selective and equally-preferring cells (**Figure 2D, left**). Second, within a given region, mean firing rates were similar across gesture types: a 2-way ANOVA showed no main effect of gesture-type on mean firing rates (F(2,1958) = 1.3899, p = 0.1591) and no interaction between gesture-type and region (F(6,1958) = 0.531, p = 0.785). Overall activity, however, did differ significantly between regions (F(3,1958) = 39.52, p = 8.7 x 10^−25^). Thus, while face-motor regions differed in overall activity, none was more active during one type of facial gesture than another.

Finally, we tested for systematic differences in the preferred gesture among highly selective cells only, in case such cells in the cingulate preferred for example, socio-emotional gestures and those in the lateral cortex, volitional gestures. We examined the preferred gesture within the subset of gesture-selective cells (defined as preference index greater than or equal to 0.5). **Figure 2D, right panel** depicts the distributions of cell selectivity preferences per region; importantly, no region contained an overrepresentation of cells selective for one particular gesture. In other words, even highly selective cells in medial (M3) and lateral (M1, PMv) regions were equally likely to prefer socio-emotional or volitional gestures. All regions contained comparable mixtures of both broadly tuned and gesture-selective neurons, and thus did not abide by a strict division of labor by gesture type as predicted by the dominant neuropsychological schema.

### Facial kinematics are encoded in low-dimensional dynamics predominantly in M1 and S1

As all face motor regions contain neurons projecting directly to the facial nucleus^9^ (with the exception of S1), it is plausible they all encode kinematics of gestures. Universal encoding of kinematics could explain both narrow tuning (a given muscle may be uniquely involved in one gesture, ie., **Figure 1E**, component 4 during threat) as well as broad tuning (the same muscle may be engaged similarly across gestures, ie., **Fig 1E**, component 2). To determine the relationship between neural population activity and facial movement, we used preferential subspace identification (PSID). PSID is a targeted dimensionality reduction algorithm that identifies and models neural dynamics specifically relevant to continuous behavior^41^. We used PSID to quantify the degree to which neural activity in each face motor region could predict facial kinematics (as estimated by marker positions) during gesture production.

**Figure 3A** depicts the decoding performance of region-specific models trained to predict the top five facial movement components during lipsmacking from behaviorally-relevant neural dynamics (see *Methods*). In brief, for each region, the method jointly considers population activity and facial movement components, in order to learn low dimensional neural dynamics driving behavior. Once this was accomplished in the training stage, each region-specific model operated on unseen neural data to predict behavior. Decoder performance, defined as the correlation coefficient between actual (**Figure 3B**, black traces) and predicted (blue traces) facial motion components on individual trials, was estimated by stratified fourfold cross-validation for each model independently. M1 and S1 consistently predicted facial kinematics better than premotor or cingulate cortex. In each region, population activity predicted facial kinematics on a trial-by-trial basis above chance (as defined by a null model in which behavioral time series were permuted and the model refit, see *Methods*). Thus kinematic information is encoded via low dimensional dynamics across face motor cortex, more so in M1 and S1 than in PMV and M3, consistent with a role for these regions in moment-by-moment, fine-grain control. Importantly, encoding of kinematic data could not explain the overlapping response properties found throughout the motor system of the face.

**Figure 3:**
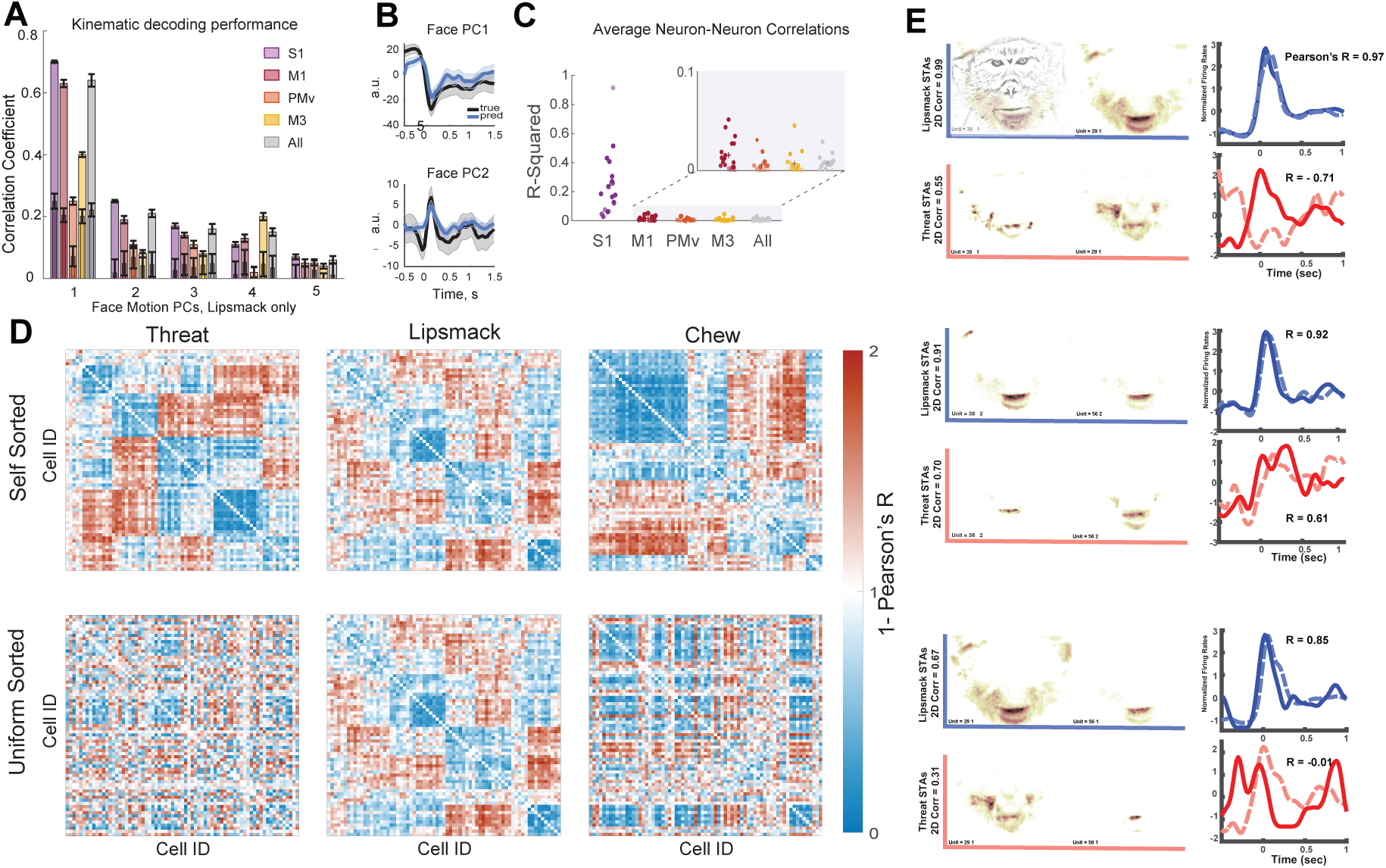
Kinematic decoding and non-preserved neural correlations. A. Distribution of kinematic information across cortical face motor areas. We used preferential subspace identification (PSID) to predict continuous facial kinematics from neural population activity. Facial movement components estimated using marker positions, (as in Figure 1E, over a different timespan of −500 to +1500 ms) were binned at 50 ms and smoothed with a 50 ms kernel. Spike data was prepared similarly. Performance of region-specific decoding models trained to predict the top five facial movement components during lipsmacking from behaviorally-relevant neural dynamics are shown. Decoder performance defined as the correlation coefficient between actual and predicted facial motion components on individual held-out trials, was estimated by stratified fourfold cross-validation for each model independently. For each model, all hyper-parameters including optimal number of behaviorally-relevant states were selected using an eightfold inner cross-validation procedure within the training data (see Methods). A null distribution was created in which the behavioral timeseries was randomly permuted, the model refit, and correlation coefficient recomputed (see *Methods*). Bar graph represents the mean correlation coefficients, which were calculated by averaging the decoding performance of twenty five pseudopopulations per cortical region (see Methods for full details) drawn from the full dataset of all available cells. Error bars represent +/- 2 standard deviations (across iterations, each first averaged across 4 outer folds). Overlaid are chance level decodings performed on a null distribution of permuted trials. B. True (black) and predicted (light blue) continuous movement of the first two facial movement components on a held out trial. Shaded error bars represent +/- 2 standard deviations (for a given iteration, calculated across held-out trials). C. For each region separately we calculated the R^2^ of pairwise neural correlations, and then averaged across all three comparisons (threat v. lipsmack, threat v. chew, lipsmack v. chew). Each dot represents the R^2^ for one recording day, for one region; opacity indicates significance level (p-value). Hashmarks indicate per region averages across recordings. Correlation values were highly dispersed, indicating the relationship between a pair of neurons during one gesture did not predict their relationship during another. With the exception of S1, R^2^ values in all regions and all recordings never exceeded 0.1 (insert), indicating neural correlations within each region were not preserved across gestures. D. Unique neural correlational structures exist for facial gestures. Pairwise correlations, 1 – R (Pearson’s correlation coefficient), between all neurons of all regions during the perimovement period of one type of gesture. Each column plots the correlation matrix for one gesture, indicated at top (threat: left, lipsmack: center, chew: right). Each element in each matrix indicates the degree to which the trial-averaged response pattern was dissimilar for the two neurons during that facial gesture. Blue indicates groups of highly correlated neurons, red indicates anti-correlations. Data analyzed from −500 to +1000 ms; spikes were binned at 20 ms, smoothed by 50 ms kernel. Bottom panel: ordering of neurons is the same for all matrices and was determined via hierarchical clustering of the lipsmacking matrix. Top panel: ordering of cells is unique for each matrix and was determined via hierarchical clustering of each facial gesture matrix separately. One typical day of recording is shown. E. Firing rates of three neuron pairs during lipsmack (blue pair of traces), and during threat (red pair of traces). Each horizontal panel depicts a pair of neurons. For each neuron, its spike-triggered movement average map is also shown. Each STMA represents the regions where movement was significantly modulated by the neuron’s firing (blue axes, during lipsmacks; red axes, during threats), when compared to a shuffled null distribution; determination of these maps was made via an unbiased, reverse correlational method (see *Methods: STMA*, for full details). For example, spiking activity of neuron 29-1 increased movement across the mouth, snout, cheeks, and ears.

### Neuron-neuron correlations are gesture-specific

How then are these gesture-specific kinematics implemented? In the domain of skilled limb behaviors, a recent theory contends motor cortex drives movement by recombining a limited set of neural states^42,43^. Given facial gestures share a common subset of effector activities^44,45^ (**Figures 1D-F**), we posited they may be enacted by conserved neural states whose dynamics drive shared kinematics. One implication of this hypothesis is that two neurons that share a response pattern in one gesture, will also share that response pattern in another: if one neuron alters its firing pattern, the second will change its firing in a coordinated manner. More generally, this means the structure of neuron-neuron correlations should be conserved across gesture types.

To address this question, we tested the degree to which gesture-specific cell-to-cell correlation patterns were preserved across gestures. In fact, we only rarely observed such coordinated changes. While we did identify neural pairs highly correlated for example, during lipsmacking that remained highly correlated during threats (**Figure 3E**, middle row; R=0.92, lipsmack, R=0.61, threat) it was also common for strong correlations in one gesture to disappear in another (bottom row; R=0.85 lipsmack, R=-0.01 threat) or even for correlations to invert (top row; R= 0.97, lipsmack, R= −0.71, threat). Even when two neurons drove movement in overlapping facial zones, during two gestures in which those zones moved similarly, the neural pairs’ activity changes were not necessarily conserved across gestures (**Figure 3E**, top row).

Next, we systematically tested for correlation preservation across gestures in the entire neural population. We computed the distance, 1 – R (Pearson’s correlation coefficient), between a cell pairs’ activity over time during the perimovement period of one type of gesture, for all neural pairs. Correlation values were highly dispersed (across all regions, all subjects, mean R^2^ = 0.0082 (threat v. lipsmack), 0.0053 (threat v. chew), 0.0092 (lipsmack v. chew). Correlation coefficients for each cortical region separately, are shown in **Figure 3C**. With the exception of S1, values in all regions and all recordings never exceeded 0.1 (insert), indicating neural correlations during one gesture did not predict those during another gesture.

Resulting matrices were ordered by inter-row similarity, revealing their rich correlational structure (**Figure 3D; top row**); each facial gesture generated a specific structure. To test if these structure were conserved across gestures, we applied the same neural ordering to all matrices (**Figure 3D**; bottom row), revealing that the correlation structure was markedly different across gestures: This could be a trivial consequence of a simpler phenomenon: if three non-overlapping populations fire exclusively during a different gesture. However, the broad distribution of selectivity indices in all regions (**Figure 2D**) directly refutes this. Facial gestures were thus not encoded via a recombination of common neural states. Instead, taken together these results imply gesture-specific subspaces of neural activity during the peri-movement epoch.

### Facial gestures are encoded prior to movement onset

As shown above, facial gestures are distinct and all cortical regions contain cells sensitive to all gesture-types. If these regions in fact encode gestures, each should contain discriminable neural representations. We found that gestures were indeed readily distinguishable from one another on the basis of their neural activity. We first examined dominant patterns of neural population activity over the entire peri-movement period. **Figure 4A** depicts clustering of individual trials by gesture-type; each trial is represented as a point in state space; axes represent the first three principal components of population spiking activity across all cortical regions (comprising roughly 30% of total variance). Within this low-dimensional space, trials clustered tightly according to gesture type.

**Figure 4:**
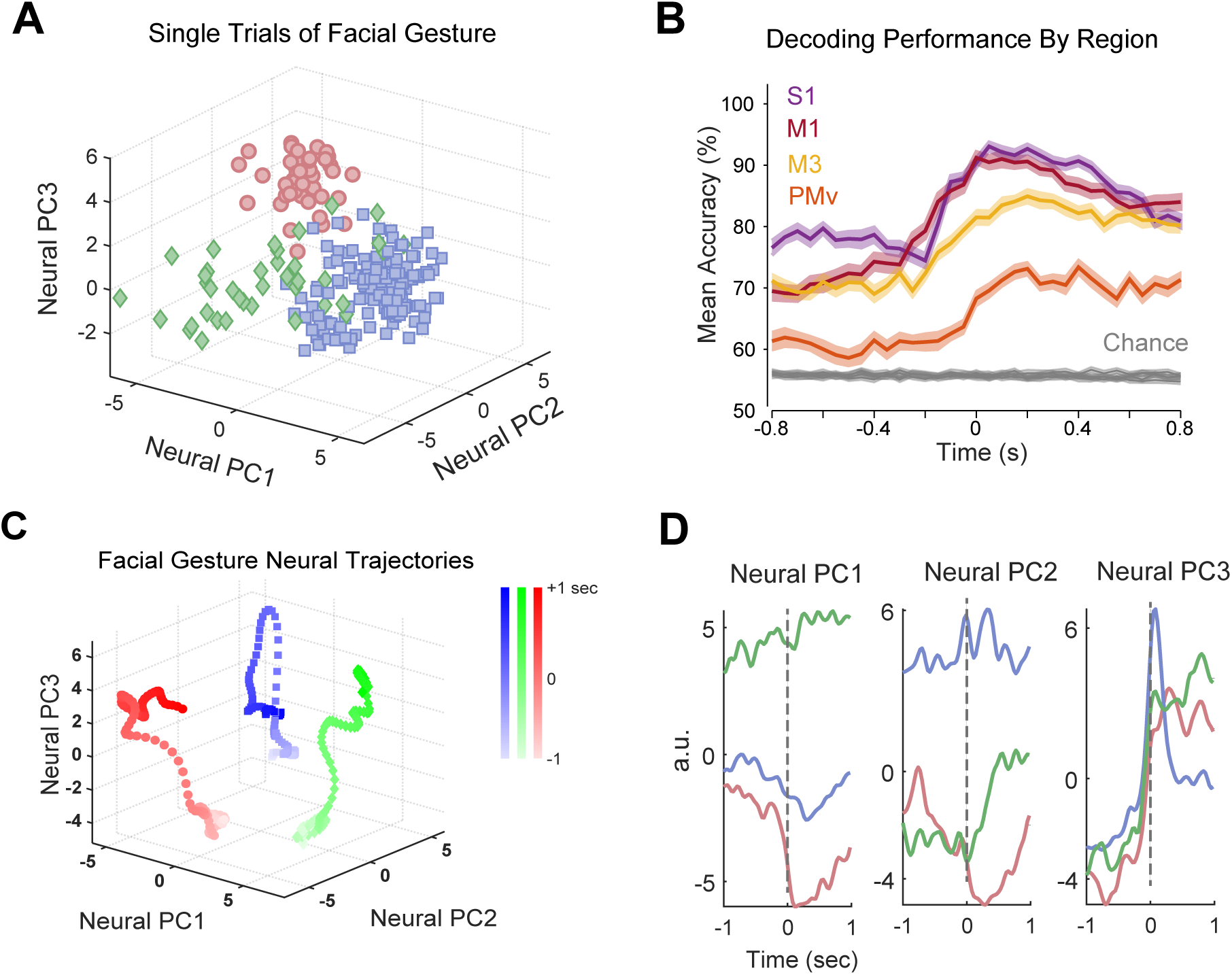
Population encoding of Facial Gestures. A. Behavioral clustering of individual trials in the neural population during naturalistic facial expression production, as depicted by PCA. Each point represents a trial (−500 ms prior to, to +1000 ms after movement onset) in 3-dimensional neural state space, colored by the facial gesture type. The top three dimensions capturing 26% (monkey B; 104 cells total) of neural variance are shown. Note that here, cells from M1, PMv, and cingulate M3 are all included together, from a single day of recording. B. Linear decoding of gesture type from population activity. Using a stratified cross validated linear SVM, we decoded categorical facial gesture from region-specific population spiking activity on a trial-by-trial basis, over 400 ms windows in 50 ms steps. Decoder performance (accuracy) was estimated by stratified 8-fold cross validation. The folds always referred to the same trials in each window throughout the trial. The mean decoding accuracy in a given time bin was calculated by averaging the decoding performance of fifty pseudopopulations per cortical region (see Methods for full details) drawn from the full dataset of all available cells. Error bars represent +/- 2 SEMs across pseudopopulations. The grey curve indicates significance threshold based on the 98th percentile of one hundred permuted decoding accuracies. Note this chance level is not ∼33% as the three gestures occurred at unequal frequencies; we chose to use all data, permute labels and repeat decoding, rather than sub-select trials. C. Facial gestures trace distinct, non-overlapping trajectories in neural space. Each trajectory is the projection of trial-averaged neural activity during one gesture type onto the top three PCs of facial gesture production. Data from −1000 ms to +1000 ms around movement onset was binned at 20 ms, smoothed with a 50 ms kernel, and very low firing rate neurons removed. Color map indicates gesture-type, and color scale indicates time during trial. The top three dimensions capturing 58% (monkey B; 103 cells total) of neural variance are shown. Note that here, cells from M1, PMv, and cingulate M3 are all included together from a single day of recording. D. Evolution of top three time-varying neural states during facial gesture production. Dashed vertical grey line indicates movement onset. Color map same as C.

To formally compare decodability across regions, we built classifier models to predict gesture type from region-specific population activity on a per-trial basis in discrete time steps (**Figure 4B**). Gesture type was decodable during the movement period in each area (peak: M1: 91%, S1: 93%, PMv: 73%, M3: 85%). To allow for fair comparisons, we balanced cell numbers across regions (N = 50 cells per region per iteration, repeated for 50 iterations; see *Methods*), though similar qualitative results (albeit with higher accuracies) were obtained using all available cells simultaneously. The time course of decodability also differed between regions: while M1 and S1 achieved similar peak accuracy, decoding performance rose and peaked earlier in M1 (91% at +0 ms) than in S1 (93% at +50 ms). Compared to M1 and S1, decodability in PMv and M3 rose more gradually (73% at +200 ms, 85% at +200 ms, respectively). Chance level was determined by one hundred permutations of trial gesture-labels and recomputation of decoding accuracies in each time bin (see *Methods*). While the precise timing of maximal discriminability varied, all cortical regions reliably encoded categorical facial gesture during movement.

Surprisingly, gesture type was not only decodable immediately prior to and during movement, but even eight-hundred milliseconds prior to gesture onset (**Figure 4B**), a window that did not overlap with overt gesture production (based on criteria for trial selection, see *Methods*). This was true not only in PMv and M3, but also in M1 and S1. To trace the temporal evolution of this discriminability, we considered neural activity during each gesture as a trajectory through high-dimensional space where each axis represents a unique time-varying neural state^46^ and used PCA to identify a low-dimensional subspace which adequately contained these data. These neural trajectories, found using the smoothed trial-averaged data, captured a large fraction of variance during gesture production (roughly 55% neural variance contained in top 3 PCs, **Figure 4C**). Gesture-specific neural states were separable long before movement started.

Notably, trajectories of different gestures never overlapped, even early in the premovement period, though gestures were requisitely preceded by stillness and began from a similar “postural” baseline (**Figure 1B**). Subsequently during movement, each gesture trajectory inhabited its own partition of state space where it traced out unique dynamics: threat and chew trajectories were linear, moving away from their origins and one another over time, whereas lipsmack displayed a rotational geometry (**Figure 4C**). Examination of the time-varying components for each neural trajectory (**Figure 4D)** revealed that most contained dynamic changes aligned to the movement, and/or sustained activity differences between gestures. Together, these results indicate that in the cortical face motor network, different facial gestures are accompanied by distinct patterns of both sustained and dynamic neural activity, both during and prior to movement.

If neural activity during facial gesture production reflects sustained state differences yoked to gesture type, it may be supported by neurons whose gesture-type selectivity is stable over time. As noted above, for a subset of cells (ie., **Figure 2C**, Unit 4) the major feature was *not* a movement-aligned activity change, but gesture-specific tonic firing, beginning prior to gesture onset and sustained throughout the movement period.

As previously shown (**Figure 2E**), most cells’ modulated their firing rate by gesture-type. A smaller but still significant fraction were modulated by time, a feature which varied by region: **Figure 2E, right** shows a significant main effect of region on the fraction of cells modulated by time (S1: 68%, M1: 47%, PMv: 24%, M3: 14%; *X*2 (3, *N* = 652) = 84.01, *p*=4.23×10^−18^); post-hoc comparisons (Bonferroni-corrected) revealed significant differences in the fractions of time-modulated cells for all inter-areal comparisons. Thus, all regions contained cells whose firing rates are relatively invariant to time, but greater fractions of cells in M1 and S1 were temporally sensitive, whereas higher-order regions M3 and PMv contained more cells with temporally stable firing rates.

### Stable and dynamic neural coding dominate distinct cortical regions during facial gestures

Though face motor cortex encodes gestures in both premovement and movement epochs (**Figure 4B, 4C**), gesture-specific activity patterns (**Figure 4C, 4D**) could be stationary or vary across time. If a region is performing distinct computations over time, the underlying neural code may differ. For example, gesture kinematics are primarily encoded in M1/S1 in the movement epoch (**Figure 3A**), yet these regions also encode gesture in the early pre-movement epoch (**Figure 4B**). Thus far, it is unclear if 1) this represents stable vs. dynamic neural representations of gesture and 2) if this varies by region. Importantly, region-specific temporal profiles of activity could reflect distinct cortical coding strategies.

To assess the stability of neural coding across time, we extended our classification analysis to cross-temporal generalization. Specifically, for each region we trained a classifier at each time point and tested against data from another time point; by systematically training and testing on all timepoints, we generated a matrix (**Figure 5, left**) in which each row corresponds to the timepoint at which the classifier was trained and the columns, to timepoints tested^47,48^. Generalization is evidence of stability as the code that differentiates behavior at one time point is functionally equivalent to that at another. Specifically here, a classifier trained on pre-movement time points able to predict behavior in the later movement period, would represent a stable coding scheme. Conversely, failure to generalize provides evidence for a temporally-local or dynamic code^49–52^.

**Figure 5:**
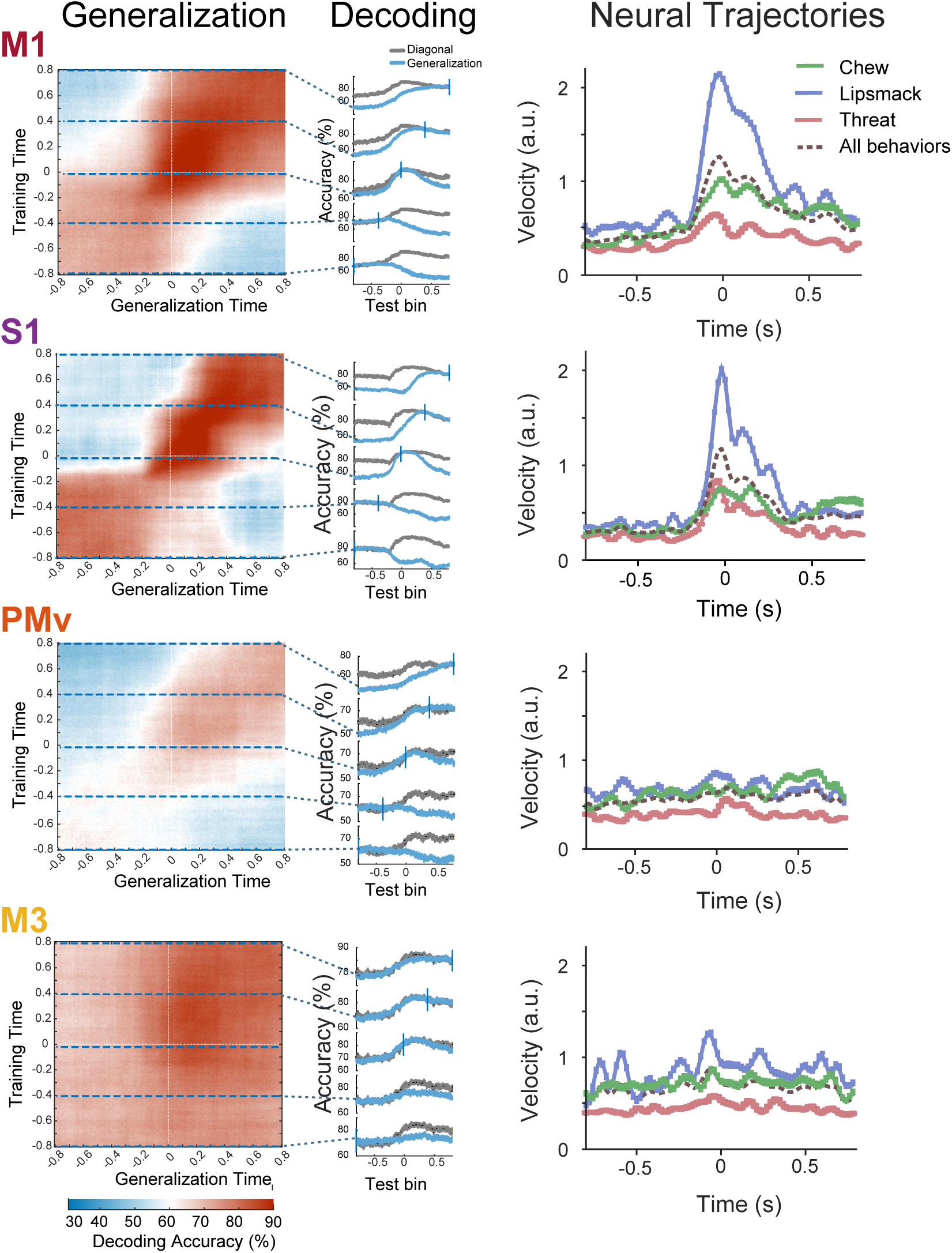
Stable and dynamic coding of facial gestures across cortex. Left panel, Full cross-temporal generalization matrices per region. Warm colors indicate above-chance decoding performance, cool colors indicate below-chance performance, white indicates chance levels. Classifiers were trained to discriminate categorical facial gesture at each 400 milliseconds time step (with 10 ms overlap) from −1000 milliseconds prior, to +1000 milliseconds after movement onset (same as Figure 4B, over longer time period). Decoder performance was then tested throughout the entire trial period. Movement onset is indicated with vertical dashed white line. Cross-validation performed as in linear SVM classifier. Matrices represent the average cross-temporal classification scores, computed separately for each pseudopopulation. Middle panel, Decoding accuracy as a function of time. Each subplot compares the performance of the decoder trained/tested on time-equivalent bins (diagonal through the matrix) (black trace), and the performance of a decoder trained on one time bin, and tested against all others (i.e., a horizontal slice through the generalization matrix) (blue trace). Where the black and blue curves overlap indicate generalization or stability of coding. Error bars represent +/-2 SEMs across iterations. Right panel, face gesture neural trajectory velocities differ across regions with dynamic and stable coding regimes. Velocity (rate of change) of facial gesture trajectories in the neural space defined by the top 12 principle components of neural activity in each cortical region. The velocity for each gesture is plotted (same color map), as is the all-gesture average (in dashed black line). Population trajectory velocity was obtained by 1) calculating the neural trajectories for each facial gesture in each region (as in Figure 5B), binned at 20 ms smoothed with 50 ms kernel 2) calculating the distance between two successive time bins in N-dimensional neural space (12-D data shown; see Supplemental for additional dimensions). Areas with stable coding also have neural trajectories with overall slower velocities, while areas with more dynamic coding have trajectories with faster velocities; periods of rapid velocity increase generally coincide with periods of dynamic coding.

We found a range of stable and dynamic coding across face motor cortex. Strikingly, area M3 in cingulate cortex displayed a highly stable code for facial gestures that persisted from 1000 milliseconds prior to, to 600 milliseconds after movement onset, visible as a solid square of high decoding accuracy (**Figure 5, M3**). Moreover, there was directional asymmetry in decoding in the cingulate: generalization performance improved when training on earlier time-points and testing against later ones (as opposed to the reverse).

Conversely, M1 housed a more dynamic, temporally-local code (**Figure 5, M1)**. Decoders trained on the premovement epoch generalized well within this time range, but did not generalize well to the movement epoch, and vice versa. The sharp transition was surprising given epochs were not experimentally defined and separated by only tens of milliseconds. S1 exhibited the most dynamic code, with highest accuracy along the diagonal. Multiple codes in S1 were detected during the movement epoch itself, visible as diagonally adjacent squares. More generally, while both regions encode gesture in pre-movement and movement epochs, this is achieved via the joint contribution of multiple temporally-distinct neural representations, which unfold sequentially within the same population. In premotor cortex, the code was neither clearly dynamic nor stable (**Figure 5, PMv**): a representation lasting roughly 400 milliseconds was detectable at movement onset and tiled the remainder of the movement epoch, but accuracy was largely confined to the diagonal. Such a pattern may indicate mixed coding schemes^50^. These findings demonstrate the different cortical face motor regions exhibit distinct temporal coding strategies.

These regional differences in temporal coding strategy were also reflected in how quickly or slowly neural states evolved over time. We estimated the instantaneous velocity of region-specific neural trajectories as the distance between two adjacent time bins in state space as a function of time; multidimensional velocity is a more sensitive measure of network state changes than global activity level^53^. In M1 and S1, neural trajectory velocities increased 200 milliseconds before movement onset, peaked approximately at onset, and then briefly decreased before increasing again at +150 milliseconds (**Figure 5, right**). By contrast, in M3 and PMv velocities were slower and constant. Largely, periods of low velocity coincided with epochs of stable coding, whereas velocity increases coincided with epochs of dynamic coding, in agreement with our cross-temporal analysis. These results demonstrate that though all cortical regions encoded all facial gesture types, each did so with a distinct temporal coding mechanism during social communication.

## DISCUSSION

Facial movements are foundational to primate communication. This form of social signaling requires the dynamic arrangement of dozens of fine muscles^1^. In this first systematic study of the neural mechanisms of facial movement control, we leveraged a new experimental approach combining three critical elements: functional targeting of cortical circuitry, multi-channel parallel recordings, and naturalistic behavior. Specifically, we adopted an approach previously used in sensory systems^54,55^, which relies on fMRI to localize regions of high functional specificity for subsequent targeting for electrophysiological recordings. This allowed us to record simultaneously from multiple areas, all specialized for control of facial movements, and thus to compare their function on the exact same trials. We deliberately did not train subjects on a specific task, as is common when studying motor physiology, but presented socially-engaging stimuli that invited naturalistic behavior. This approach is critical for studying facial gestures, a behavior that is strongly driven by social and cognitive cues such as context, inner state, and the relationship between individuals.

Despite the complexity of facial musculature, facial gestures evoke highly stereotyped patterns. These patterns are repeated throughout life^56^ and involuntarily^57^, suggesting reliance on subcortical mechanisms, potentially via dedicated central pattern generators^26,58^. Contrary to expectation, we found that (i) cortex *is* intimately involved in facial gestures, that (ii) it is broadly and equally involved in distinct motor behaviors including socio-emotional expressions and voluntary movements, and that (iii) in fact multiple cortical areas are involved. This distributed network with both lower-order (M1/S1) and higher-order (PMv/M3) areas projecting directly to the facial nucleus^9^, necessitates an organizational principle for generating coherent motor outputs. We found that although neurons throughout the system showed mixed selectivity to facial movements, different facial gestures were enacted by unique temporal patterns of activity. Critically, neural representations were encoded via regionally-distinct temporal mechanisms, forming a continuum of stability in the code for facial gestures across cortex. Naturalistic facial communication is not strictly manifested in the coding complexity of single cells, but instead represented throughout the cortical face motor system, where hierarchy is in the temporal domain.

### Double dissociation of medial and lateral face motor regions based on behavior type is not supported by single cell activity

Following a well-accepted neuropsychological schema of cortical face motor control^32,59–61^ partially supported by neuroimaging studies^62–68^, we expected an enrichment of cells tuned to socio-emotional gestures in the medial cingulate cortex, and to volitional movements (ie., chew) in lateral motor and premotor cortices. Unexpectedly, there were no differences in quantity or gesture preference of selective cells between medial and lateral cortices (**Figure 2D**). Such broad tuning of single cells resembles that in a different effector system, not during spontaneous behavior, but after extensive training: simple upper limb movements are accompanied by similarly broad tuning curves. This may reflect a principal role for motor cortex in generating temporally-patterned outputs, in lieu of representing overt behavioral categories^69–71^.

Our results refute the neuropsychological schema of facial movement control, and instead reveal facial gesture production as a distributed process dependent on multiple cortical regions. Careful past work suggested this: functional neuroimaging in the macaque supports qualitative, rather than quantitative differences between medial and lateral face motor areas with respect to behavioral category^68^. Moreover, a systemic radiological study of 211 patients with isolated cortical lesions and facial paresis found only four with emotional facial palsy, all of whom showed concurrent impairment in volitional movements^72^. How then can the apparent double-dissociation of facial movements after lesioning be explained? Our results suggest a distinct etiology. Here we show that the medial cingulate face motor region (M3) houses abstract and stable representations over long timescales. We propose that the cingulate cortex is required not to drive a unique set of facial movements, but to represent contextual information and provide it to lateral face motor regions. In support of this hypothesis, there lies another region just anterior to the face motor region M3, which is not correlated with facial movement but social context^68^. Under this hypothesis, the dearth of internally-generated facial gestures seen in medial lesion patients^59^ as well as those with akinetic mutism^73,74^, can be seen as an effective disconnection. More generally, the apparent discrepancy between 1) non-specific neural activity and 2) seemingly specific behavioral impairments after lesioning of the same site, may be explained by single cell activity carrying multiple messages and in itself may not be easily related to the contribution of neural regions to motor output^75^.

### Context-dependent encoding of facial kinematics

We found that the encoding of kinematic information could not sufficiently explain the response properties of neurons throughout the motor system of the face. Kinematic information was decodable across cortex but concentrated in primary motor and somatosensory cortices (**Figure 3A**). Additionally, facial gestures were encoded in low-dimensional neural dynamics (**Figure 3A, 4C**), consistent with the neural manifold theory^43^. One relevant extension of this theory is that distinct skilled limb behaviors are generated by recombining a shared set of correlated neural activity patterns, preserved across behaviors^42^. By contrast, neural correlation structure was not preserved across gestures (**Figure 3D**) and was gesture-specific. The encoding of facial gestures follows a distinct logic from that of the upper limb motor system, at least in paradigms following extensive training. Furthermore, our findings imply the existence of gesture-specific subspaces of neural activity, an increasingly recognized computational mechanism by which cortex may organize and segregate interacting information streams^76–80^. Together these results indicate cortical regions are not merely coding facial kinematics at any moment, but doing so in a context-dependent manner, related to the behavioral state in which they were embedded. Context-sensitivity may be driven in part by cells that were not explicitly movement-related (e.g, **Figure 2C**, Unit 4) but modulated their baseline rate in a context-dependent manner^81^.

Similar impact of context on neural coding has been reported in sensorimotor cortex, attributed to a representation for phoneme coarticulation during speech^82,83^. In the limb, context-dependent coding has be found in mirror neurons^84^, and is modulated by learning and related to the degree to which behavior is naturalistic of restricted^85,86^. During naturalistic facial gestures, context-dependent encoding could efficiently account for individual muscle movements influenced by surrounding kinematics and/or meta-variables of the environment from which the gesture is generated. The gestures we studied are related to dramatically different behavioral contexts and could therefore be enacted by unshared neural states. Clinically, the context-dependency we show here may have important implications as using such signals may improve decodability of more complex, naturalistic behaviors with brain-computer interfaces in patients, a current challenge in the field^87,88^.

### Neural states segregate long before movement onset

The segregation of neural activity into distinct sub-spaces did not occur just prior to, or at movement onset, but long before analogous behavioral trajectories separated from a common, neutral basis (**Figs. 1F and 4C**). In the motor system of the upper limb, preparatory activity is an essential epoch in which neural computations (restricted to a null-output space^77,79^) bring the system to an initial state from which movement subsequently unfolds^68–71^. It is possible that in the facial motor system, preparatory activity monitors social context which changes on a slow timescale. This may explain why pre-movement neural trajectories of different facial movements never overlapped. However, preparatory activity also had unique temporal properties in different cortical regions.

In M1, preparatory activity did not generalize from pre-movement into the movement epoch. If preparatory activity in M1 functioned as a sub-threshold form of movement, we expect a code generalizable across epochs^89–91^, not unlike that seen in the cingulate (**Figure 5**, bottom panel). Instead, a sharp transition at facial gesture onset (**Figure 5, top panel**) corresponded with a rapid rearrangement of activity in neural state space in M1, reminiscent of transitions from preparatory (output-null) to movement-related activity (output-potent) in studies of the upper limb. This theory has relied heavily on highly trained reaches^92–95^. Our results demonstrate for the first time that preparatory activity in M1 may play an analogous role in the face motor system, and during social behaviors more generally.

Preparatory activity in the medial cingulate (M3) was also predictive of upcoming facial gesture well before it occurred. Critically, the M3 code generalized from premovement throughout the entire movement period. Corresponding, M3 contained more gesture-selective cells with temporally stable firing rates (**Figure 2E**), and slower trajectory velocities (**Figure 5, bottom right**). It is not clear however, what mechanism prevents the stable M3 code from producing premature gestures during the anticipatory period. It is possible that the net effect of this area on muscles is either too weak or gated by downstream inhibitory mechanisms^96,97^. Temporal stability of neural coding in PMv fell between the extremes of M3 and M1/S1. Relatively little kinematic information was present in M3 or PMv (**Figure 3A**), signifying their neurons’ tuning properties cannot be explained by kinematic sensitivity alone, despite the fact that decoding accuracy was comparable (M3) or only slightly lower (PMv) to that of regions in which kinematic information was plentiful (e.g, M1). It is likely that parts of preparatory activity are related to variables other than context, that are differentially processed along the motor hierarchy. Future study is necessary to identify these variables and their contribution to social interactions.

### A hierarchy in the temporal domain

Sensory areas in cortex are hierarchically organized such that coding complexity monotonically increases from low to high regions (e.g., from orientation tuning in V1 to face selectivity in IT). In contrast, hierarchy in the motor system is poorly defined. High level motor areas gain direct access to motoneurons via anatomical connections^98,99^, an arrangement inconsistent with a strict step-wise hierarchical framework. Modular organization of function, investigated in specific systems such as reach vs. grasp, or posture vs. reach, has been challenged by studies showing colocalization of coding neurons in single cortical areas^100,101^. Subsequent frameworks of the motor system have emphasized task-oriented optimal control^102,103^ or task-specific temporal dynamics occurring at different regions of the motor cortex, even though single cells are often similarly sensitive to task variables.

Our results confirm and expand these concepts by demonstrating that informative neural states present throughout the face motor system exhibited distinct, regionally-specific temporal dynamics (**Figure 5**). Across cortex, these dynamics form a novel continuum of coding stability, from long (cingulate) to medium (premotor) to short (primary motor and somatosensory), with lower-order regions using correspondingly shorter timescales appropriate for movement generation.

Anatomic segregation of dynamic and stable coding, thought to reflect specialization in the type of information a region processes, has been reported in prefrontal and sensory cortices during working memory tasks^104,105^. Experimental and theoretical work has shown primate cortex houses a hierarchy of intrinsic timescales which may reflect areal specialization for task-relevant computations, with sensory regions showing shorter timescales, and high-order or “transmodal” regions including the anterior cingulate, consistently showing longer timescales^106–109^. We demonstrate a similar hierarchy of timescales across cortical motor regions supports gesture production during naturalistic social interactions. What advantages might a network that houses temporally-sensitive and insensitive forms of the same motor information, have over alternative arrangements?

### A novel functional division of labor in the cortical face motor system

We purport that while all regions represent socio-emotional and volitional facial gestures, the primary motor and somatosensory cortices do so with fast dynamics driving fine-grain kinematics and real-time adjustments, while the cingulate does so with highly stable representations. During gesture production, temporally-local representations in M1 and S1 underwent rapid reconfigurations. By definition such dynamic representations encode temporal information critical to organizing both internally-generated and externally-cued motor behaviors. Appropriately both regions contained substantive kinematic information, consistent with a role for controlling movement, in accordance with work showing cued articulatory movements and speech can be decoded from human sensorimotor cortex^110–112^.

The precise functional roles played by the slower evolving dynamics in premotor cortex, and strikingly stable dynamics in the cingulate, are still unclear but may be hinted at by anatomy. Anterior cingulate subregions containing projections to facial nucleus^9^, spinal cord^113,114^, and motor cortex^98,115^ receive afferent inputs from prefrontal cortex and limbic structures^116–118^, enabling linkage of contextual information to subsequent action plans^118–122^. This positions M3 as a potential interface by which contextual information operating on longer timescales can impact facial movements. It is at this very interface we find a stable code for facial gestures, which allows information to be held and then extracted via a constant decoding scheme whenever needed.

Successful sociality requires coordination over several distinct timescales. Our findings necessitate the proposal of a new functional division of labor amongst cortical face motor regions which serve this purpose. In facial gesture production, sub-second kinematics are the end product of a larger sensory-motor process wherein cognitive variables like visual information^55,123^, internal states^124–126^, and somatosensory feedback each play a role^127^. This continuum of temporal stability in the neural codes for facial gestures could provide a mechanism by which a motor system heavily reliant on contextual cues, is able to integrate these cues while producing well-timed, accurate, and interpretable motor outputs. We hypothesize the regionally-specific temporal stabilities reflect a broader mechanism by which a cortical network impacts social behavior over multiple distinct timescales.

### Data availability

The datasets generated and analyzed during the current study are available from the corresponding author on reasonable request.

## Author contributions

GRI, YV, YP, and WF designed the study. GRI and YV acquired the data. GRI and YV analyzed the neuroimaging data. GRI analyzed the electrophysiology data here. AG and MS provided surgical expertise. All authors assisted on data interpretation. GRI, YP, and WF wrote the paper.

## ACKNOWLEGEMENTS

We thank the staff at the Cornell Citigroup Biomedical Imaging Center for assistance with data acquisition, A. Gonzalez for help with animal training and care, veterinary services and animal husbandry staff of The Rockefeller University for care of the subjects, and L. Yin for administrative assistance. We thank H. McAdams for thoughtful discussions. We are especially grateful to S. Shepherd for his work on the imaging analysis of subject 2, to S. Schaffelhofer for his development of neurosurgical tracking technologies (now cortEXplore, GmbH, Linz Austria), and to both S. Shepherd and S. Schaffelhofer for neurosurgical targeting of subject 2.

## METHODS

### Subjects

Two pair-housed male rhesus monkeys (subject 1: Macaca fascicularis, age 5 years; subject 2: Macaca mulatta, age 7 years) participated in the study. Procedures conformed to applicable regulations and to NIH guidelines per the NIH Guide for Care and Use of Laboratory Animals. All experiments were performed with the approval of the Institutional Animal Care and Use Committees of The Rockefeller University and Weill Cornell Medical College. All subjects were fitted with head restraint prostheses using standard lab approaches (Fisher and Freiwald, 2015).

### fMRI, Task and Behavior

Of note, Subject 2 was previously fMRI-mapped as fully described in Shepherd & Freiwald, 2018. Subject 1 was fMRI-mapped in an analogous fashion, outlined fully here for clarity. Prior to scanning, monkeys were trained to sit in a sphinx position and maintain fixation on a red dot at the center of the screen for 2 - 4 s to receive fluid reward while blocks of pictures or videos were presented, interleaved with baseline periods during which only a fixation dot was present. During imaging, subjects sat in an MRI-compatible NHP chair with the head fixed at isocenter. Gaze position was monitored at 120 Hz with an MRI-compatible infrared eye-tracker, and calibrated at the beginning of each scanning session. All behavioral and stimulus display parameters were under control of Presentation software (Neurobehavioral Systems). During functional scans, stimuli were video-projected at 60 Hz resolution on a back-projection screen placed 35 cm in front of the subjects’ eyes.

All subjects performed the social-interaction free-viewing task as described fully in Shepherd & Freiwald, 2018. Briefly, we presented ten-second video clips of monkeys producing dynamic facial expressions facing forward, (simulating direct eye contact with the subject), while we performed whole-brain fMRI and video-recorded the subject’s face. A fixation point was present only during baseline periods; monkeys could freely move their eyes to explore the videos as long as their gaze stayed within a virtual window of the size of the frame of the video and additional 2° surrounding it. No reward was given during the social-interaction task, and the reward tube was removed to prevent occlusion of the subject’s face. In addition, each subject performed an orofacial-motor localizer in which sparse, intermittent fluid reward was delivered during an unrelated passive-fixation task.

Facial movements were captured at 15 Hz using an MR-compatible infrared video camera (MRC), whose acquisition was synchronized to the first TR of each scan run. Two independent examiners manually annotated offline the onset and category (lipsmack, threat, chew, drinking, other) of each facial expression.

### fMRI, Imaging

Each subject was previously implanted with an MR-compatible headpost (Ultem; General Electric Plastics) using zirconium oxide ceramic screws (Thomas Recording; Rogue Research), medical cement (Metabond and Palacos) and standard anesthesia, asepsis, and post-operative treatment procedures. For subject 2, imaging data was collected and analyzed as described in Shepherd & Freiwald 2018. For subject 1, imaging data was collected in a 3T Siemens Prisma scanner. Functional images were gathered with a custom-designed surface 8-channel receive radiofrequency coil and a horizontal single loop transmit coil (L. Wald, MGH, Martinos Center for Biomedical Imaging) in echoplanar imaging sequences (EPI: TR 2.25 s, TE 17 ms, flip angle 79 degrees, 1.2 mm^3^ isotropic voxels, FOV = 96 mm; matrix size = 80 x 80 x 45, horizontal interleaved slices, 2x GRAPA acceleration; the number of volumes varied per run, 85, 168, or 204). To increase the contrast-to-noise ratio (CNR) a contrast agent MoldayION (monocrystaline iron oxide nanoparticles, 6-9 mg/kg) was injected intravenously into the saphenous vein at the start of each imaging session (cite Mandeville).

Anatomical volumes were acquired in a separate session while subjects were anesthetized (ketamine and isoflurane, 8 mg/kg and 0.5-2.0%) and positioned in an MR-compatible stereotactic frame, using a custom horizontal single channel receive coil (L. Wald, MGH, Martinos Center for Biomedical Imaging). Anatomical T1-weighted images were constructed by averaging six repetitions of a T1-weighted Magnetization-Prepared Rapid Gradient Echo (MPRAGE) sequence. In addition, a head-neck CT (computed tomography) scan acquired on the same day, provided high resolution exterior anatomy.

### fMRI, Analysis

All imaging analyses were done with FreeSurfer and FS-FAST (v6.0), using custom MATLAB and Linux-shell scripts. Raw image volumes were 2D (slice-wise) motion and time corrected (AFNI, 3dAllineate, version AFNI_2011_12_21_1014), aligned to high-resolution anatomical T1, unwarped (JIP Analysis Toolbox, v3.1), smoothed (2 mm Gaussian FWHM) and masked. The first four volumes of each functional run as well as time-points having an absolute z-score intensity >3 were excluded from further analysis.

We constructed two separate models to localize cortical face motor regions in each subject: in the first, manually-annotated facial gesture onset times were convolved with the MION hemodynamic response function (HRF) to generate the primary regressor of interest (‘face-gesture map’, hereafter); in the second model, fluid-delivery times (available from Presentation log files) were treated equivalently (‘face-motor map’). Average signal intensity maps for each contrast were computed with FS-FAST function selxavg3-sess. The MION HRF was modeled as a wide gamma function with a steep initial slope (delta = 0; tau = 8000; alpha = 0.3; dt = 1 ms), and discretized to the length of the TR (2.25 s), in order to be used as a regressor in the GLM. In both models, nuisance regressors included 2^nd^ order polynomial for drift, and the top 3 head motion PCs obtained from 2D slice-wise motion correction (3DAllieniate). The resulting voxel-wise beta-weight maps were not further corrected for multiple comparisons. To identify significant voxels in both the facial-gesture and the facial-motor maps, we conducted conjunction analyses using logical ANDs, taking the least significant p-value from each contrast entered in the conjunction. This conjunction map was used in all future planning steps. This allowed us to reliably identify voxels primarily correlated with both types of orofacial movements, and not body movement which occasionally accompanied facial gestures during the social-interaction free viewing task.

### fMRI-Targeted Electrophysiology; Surgical Implantation

We used cortEXplore (GmbH, Linz Austria), an integrated functional and structural MRI-based stereotaxic planning and intra-surgical tracking software, to calculate the desired positions of 6-8 floating microelectrode arrays (MicroProbes for LifeSciences) based on our functional localization of cortical face-motor regions and contrast-enhanced, high-resolution anatomical MRIs. This allowed us to visualize blood vessels to be avoided and maximize overlap between functionally-active face motor regions and array footprint.

For each subject, all functional maps and a high-resolution CT were aligned to the anatomical MRI. All surgical planning thus occurred in the subjects’ native spaces. Target locations in somatosensory cortex (S1), primary motor cortex (M1), ventral premotor cortex (PMv), and medial cingulate cortex (M3) were identified per subject on the basis of 1) peak functional activity 2) concordance with known anatomical landmarks and stereotactic coordinates, and 3) avoidance of blood vessels. For each target location, we defined a corresponding entry point on the cortical surface; together these two points define a 3D trajectory. Because each electrode’s length in an FMA can be specified at the time of manufacture, each FMA included electrodes of varying length (+/- 0.5 mm from target) to maximize gray matter neural activity recordings.

Under sterile technique and anesthesia, we first registered the subject in CortExplore. Registration creates a spatial connection between the 3D virtual medical images space and the 3D surgical subject. This is achieved by taking serial measurements of the 3D surgical subject space with an optically tracked probe, whose configuration is known and defined in the 3D virtual space by the neuronavigation software. We registered the subject in two steps – first with seven fiducial markers on the subject’s acrylic implant, and second, with 5-8 surface registrations, in which the probe scans the surface of the surgical subject, registering it to its corresponding virtual surface. For each subject, the registration error was <200 microns.

After craniotomy and durotomy, each FMA was loaded onto the optically-tracked probe, aligned with the predefined trajectory, and advanced slowly into cortex. In all cases the 3D angular error between the actual and predefined trajectories was <1 degree.

For subject 2, six 32-channel arrays were implanted: one in somatosensory cortex (S1), two in primary motor cortex (M1), one in ventral premotor cortex (PMv), and two in medial cingulate cortex (M3). For subject 1, seven 32-channel arrays and one 16-channel array were implanted: one in somatosensory cortex (S1), two in primary motor cortex (M1), one in anterior primary motor cortex (F4), two in ventral premotor cortex (PMv), and one 32-channel and one sixteen-channel, in medial cingulate cortex (M3). All arrays were implanted in the subjects’ right hemispheres. The anterior primary motor cortex array corresponds to the ventral portion of area F4, a functionally distinct subdivision of precentral gyrus (Rapan, 2021) which has been associated with simple orofacial and tongue movements (Maranesi, 2012). The ventral premotor cortex arrays correspond to area F5; both regions house populations of mouth motor neurons responding to communicative gestures such as lipsmacking (Ferrari 2003). The medial cingulate cortex arrays correspond to the face representation in area 24c which has been extensively anatomically described elsewhere (Morecraft 2012).

Each 32-channel FMA consisted of 36 electrodes total: 28 platinum/iridium recording electrodes of varying lengths (1.2 - 2.8 mm in length for S1, M1/F1/F4, PMv/F5; 5 - 9 mm for M3/24c), 4 low impedance (10 kOhm) pure iridium electrodes for microstimulation, and 4 low impedance (<10 kOhm) reference and ground electrodes (2 and 2, respectively). Electrodes were arranged in a 9×4 triangular matrix (400 micron spacing) on a 1.8 x 4 mm ceramic chip. Each 16-channel FMA consisted of 18 electrodes total: 14 platinum/iridium recording electrodes of various lengths, 5-9 mm varying from x-y in m1, etc, 2 low impedance (10 kOhm) pure iridium electrodes for microstimulation, and 2 low impedance (<10 kOhm) reference and ground electrodes (1 and 1, respectively). Electrodes were arranged in a 4×4 triangular matrix (400 micron spacing) on a 1.95 x 2.5 mm ceramic chip. In the case of the anterior motor cortex in subject 1, we used a high-density array for spacing constraints (250 micron spacing).

### Electrophysiology; Behavior

Prior to recordings, subjects were trained to sit head-fixed in a vertical NHP chair and watch three to five second video-clips containing nature scenes (but not conspecifics) for juice reward. During experimental recordings, the reward tube was removed to prevent occlusion of the subject’s mouth and disambiguate social facial gestures from ingestive movements.

We created a number of distinct visuo-social contexts which elicited facial movements from the subject; each is described fully below. In brief, 1) movies of conspecifics 2) a monkey avatar and 3) real-life social interaction with a conspecific or human, all elicited socially-meaningful facial gestures. These visuosocial recording blocks were interleaved with blocks of non-social context, in which ingestive movements (chewing, drinking, licking) were elicited with food, and blocks of rest, in which the animal sat quietly inside the darkened recording room. A typical recording session included two to five repetitions of each block-type.

### Behavioral monitoring

During recordings, subjects’ facial movements were continuously captured (70 Hz, 1280 x 1080 pixels) with a high resolution, monochrome camera (FLIR, BFS-U3-13Y3M-C) or a Flex 13 Optitrack system (120Hz) mounted ∼ 70 cm from the subjects’ face. Videos were captured by a dedicated Windows computer running Spinview or Motive software and synchronized to the neural acquisition system via a hardware trigger (GPIO; 6-pin Hirose cable), which provided a frame-by-frame digital timestamp. A second, identical camera was used to monitor body movements.

### DeepLabCut

We used DeepLabCut (version 2.1.8.1) for two-dimensional markerless face tracking (Mathis et al, 2018, Nath et al, 2019). For each subject, we labeled 40 (subject 1) or 50 (subject 2) facial points in 185 frames from 12 videos (see **Figure 1B**). Because behavior was often sparse, the frames were chosen manually to include the broadest range of observed facial configurations. We used a MobileNetV2-1-based neural network with default parameters for 1030000 training iterations (94% of frames were used for training, 6% for testing). The training error was 1.28 pixels, the testing error was 5.46 pixels. We used a probability cutoff of 50% for all facial coordinates in future analysis. The subject-specific networks were then used to analyze all facial videos of that subject in all experimental contexts.

### Visuosocial Paradigm; Conspecific Movies and Monkey Avatar

All visual stimuli were presented and behavior controlled using MonkeyLogic (v2.2), running on a Windows computer system which sent triggers to the TDT data-acquisition system via an analog and digital input/output PCIe-6343 card. Subjects viewed visual stimuli on a 56 x 24 cm (1024 x 768 resolution) screen at 58 cm from the eyes, with a refresh rate of 60 Hz. Eye position was recorded with ISCAN (ETL-200) at 120 Hz.

A trial began with 1000 milliseconds of required fixation within an 8 degree window, after which a video stimulus appeared; the subject was able to freely move his eyes to explore the videos as long as his gaze stayed within a virtual window of the size of the frame of the video and additional 2° surrounding it. No reward was given during these runs, and the reward tube was removed.

### Conspecific Movie Stimuli

Video stimuli were recorded with a GoPro7 camera attached to monkeys’ home cages. We extracted ten-second clips of monkeys engaging in common social behaviors like grooming and playing, but also non-social behaviors like eating, drinking, and idling. Videos contained one or two monkeys, either male or female, and a subset of these monkeys were also personally familiar to the subject. To minimize behavioral adaptation, each clip was phase-scrambled (preserving low-level motion, color and luminance intensity); a single stimulus set shown during a single recording run consisted of 7-10 individual clips and their matched scrambles presented in a pseudo-random order.

### Monkey Avatar Stimuli

For a subset of recordings, subjects viewed an interactive monkey avatar, running in Unity on a separate PC, whose facial movements were controlled by the experimenter in real time.

### Visuosocial Paradigm; Real-life social interaction

A conspecific from the colony or a human experimenter was seated across from the subject; the pair were intermittently visible to one another through a partition, which was opened/closed in ∼10s segments to grossly approximate the timescale of other visuosocial contexts. Individuals with whom the subject interacted included males and females, both personally familiar and unfamiliar to the subject.

### Non-social / Ingestive Paradigm

In these blocks an experimenter seated to the subject’s left, fed the subject intermittently with fruit and dry snacks.

### Neural activity acquisition

From the implanted arrays, we recorded simultaneous neural activity from 256 (Subject 1) or 192 (Subject 2) electrodes at full bandwidth with a RZ2 Biosignal Processor (Tucker Davis Technologies). Neural activity was sampled at a rate of 24414 Hz (16-bit resolution) and stored synchronously to disk together with behavioral videos and eye movements. Raw recordings were filtered offline to obtain single-unit and multiunit activity.

### Spike sorting

All spike sorting was done offline. We first applied Waveclus^128^ to filtered, single-unit activity for automatic sorting, and then if necessary, refined manually with Plexon Offline Sorter. Sorting was performed on across all recording blocks within a day, in order to identify stable neurons with minimal drift (ie., identifiable in all recording blocks). For the purposes of this paper, waveforms identified on separate recording days were always treated as separate neurons.

### Microstimulation

In order to validate that our fMRI-localization and surgical implantation had correctly targeted face motor regions, we performed a series of microstimulation experiments in Subject 1. At each of the four low impedance electrodes on each array, we delivered a 50 ms burst of biphasic pulses at 300 Hz; for electrode, a current threshold was identified, defined as the amplitude at which 50% of deliveries elicited an obvious facial muscle twitch. After we identified threshold amplitude, we delivered 50 repetitions each of stimulation at threshold and x1.5 suprathreshold, randomly interleaved. This procedure was repeated for all low impedance electrodes on all arrays. Simultaneously recorded neural activity from arrays not undergoing stimulation, as well as facial videography was stored as previously described.

### Behavioral Scoring, Movement Onset Detection

Two independent experimenters reviewed and manually annotated the behavioral videos to obtain the onset, duration, and category of facial gesture. Each frame was scored as containing threat, lipsmack, chewing, fear grimace, yawn, nostril movements, eyebrow movement, or other. Behavior was manually annotated at the same temporal resolution as the camera capture (70 Hz for subject 1, 120 Hz for subject 2), though in subsequent analyses, we flexibly selected facial gesture events such as to only include events preceded by an appropriate amount of stillness.

### Behavioral Analysis; Facial gesture clustering and trajectories

We extracted the two-dimensional positions of forty (subject 1) or fifty (subject 2) continuously-tracked facial markers (DeepLabCut) in the perimovement period (−500 to +100 milliseconds around movement onset). Data was binned at 100 ms, smoothed with a 50 ms kernel, and PCA applied to extract the top *N* components accounting for 95% of behavioral variance. These components were then reduced to two dimensions using tSNE and plotted; in **Figure 1C**, each dot represents an individual timepoint during an occurrence of threat (red), lipsmack (blue), or chew (green), demonstrating that configurations of facial markers tended to cluster by gesture-type.

To determine the facial gesture behavioral trajectories (**Figure 1F**), we extracted the same facial marker position data from 1000 before, to +1000 milliseconds after facial gesture onset, re-binned the data at 10 ms, and smoothed with 50 ms kernel. We arranged the data into a matrix *M* ∈ ℝ ^FT x N^, where N is the number of facial marker positions, T is the number of timepoints, and *F* is the type of facial gesture. We performed PCA using singular value decomposition on the mean-centered *M*, handling each facial marker position as a variable, to obtain the *N x N* matrix of face gesture movement PCs, as well the PCA loadings of each facial marker (**Figure 1D**). The top five facial gesture principle components (**Figure 1E**) routinely accounted for >90% of behavioral variance and varied systematically across gesture type.

### Spike-triggered Movement Average Analysis

In order to identify regions of the face whose movement was significantly modulated by the firing of a neuron of interest, we constructed a spike-triggered movement average (STMA) movie for each cell^129^. Facial videos were cropped, down-sampled, and the per-pixel luminance derivative was taken as a proxy for movement (due to head-fixation, the vast majority of luminance changes represented facial movements). STMAs were computed pixel-wise:

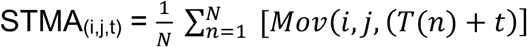

where i and j are the coordinates within the frame, t is the time from spike, Mov is the movement (luminance derivative), and T(n) is the time of the n’s spike. We examined a period up to 1 second post-spike. To determine statistical significance, we then generated a null distribution by simulating a Poisson-spiking neuron with the same average firing rate as the real neuron, and re-computing the STMA on this simulated neuron; this process was repeated 100 times. We then converted each pixel of the real STMA to a z-score by the mean and standard deviation of the null distribution STMAs. We performed minimal frame-by-frame smoothing with a 2D Gaussian kernel (size = 3 pixels). In order to be considered significant, we applied a simple threshold: a pixel needed to retain an absolute z-score of >3 in 2 or more temporally-adjacent frames (under the assumption that true spike-triggered movement fields should be relatively smooth in time).

### Common Neural Data Preprocessing

In all analyses that included examination of the premovement period, only trials of facial gestures which were preceded by adequate rest were included. This was to allow for appropriate comparison across conditions, given all facial gestures began from a stereotypical rest position. All spikes were aligned to the onset of movement (t = 0) as determined above, using the per-frame digital timestamp. In all analyses that follow, very low firing rate neurons (< 0.1 Hz) as well as those with significant drift (ie., not present throughout entire day of recording) were excluded.

### PETHs, firing rate sensitivity to behavior and/or time, selectivity indices

To assess for modulation, spike counts on each trial were aligned to hand-scored movement onset, and binned in the 500 milliseconds prior to onset, and 500 milliseconds after onset. Spike counts were entered into a 2-way ANOVA (gesture type; 3 levels: threat, lipsmack, chew; time; 2 levels: before, after onset) and significance threshold set at p < 0.05 (multiple comparison corrected with Tukey’s HSD test). We considered a cell modulated if there was a significant effect of any factor or interaction. Peri-event time histograms (PETHs) were computed by aligning spikes to gesture onsets, binning at 1 ms, and convolving with a 50 ms Gaussian kernel. We analyzed the modulation of firing rate with respect to behavior as well as time for each cell individually. To assess single cell selectivity to any given facial gesture, we computed a per-cell gesture preference index, defined as:

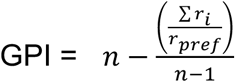

where n = number of facial gestures tested, r_i_ is the firing rate during gesture i, and r_pref_ is the maximum firing rate achieved during any gesture. This index is 0 when a cell fires equally for all gestures, and 1 when a cell fires for one gesture exclusively (Raos 2006).

### Neural Pseudopopulations Creation

For those analyses in which we quantified the relationship between neural activity and behavior on a single trial basis in order to compare across regions (∫, kinematic decoding; Figure 4B, categorical time-equivalent decoding; Figure 5, cross temporal decoding), we collapsed across recording days and subjects in the following manner: we organized the data into pseudopopulations; one pseudopopulation contained fifty cells from a given cortical region, drawn without replacement from the overall pool of available cells (across all recording days and both animals). One pseudopopulation was a raster of spike counts binned at 1 millisecond, of dimensions nTrials x nTimepoints x 50 cells. Each pseudopopulation therefore contained a unique combination of neurons and their associated behavioral trials. We created fifty (Figures 4B, Figure 5) pseudopopulations for each cortical region separately. In all cases, decoding was performed on each pseudopopulation separately, and figures represent the mean decoding performance (either accuracy or correlation coefficient), averaged across pseudopopulations. As the focus of much of our analyses was inter-cortical area comparisons, this provided normalized decoding performances per equal-sized group of neurons in each region.

### Categorical Decoding Analysis

In order to decode categorical facial gesture from neural population activity in each of the cortical regions, we used a linear multiclass support vector machine (SVM; Matlab *fitcecoc*). Fitcecoc trains an error-correcting output codes model using *K*(*K* – 1)/2 binary support vector machines (SVM) in a one-versus-one design, where *K* is the number of unique categorical labels; here, three binary classifiers were used to classify the three gestures. The input to the classifier at each time point was an nTrials × nCells matrix of spike counts, binned at 400 millisecond time windows with 50 millisecond overlap. We performed light denoising first by 1) taking the top N components (PCA) accounting for >98% of variance and re-projecting the data onto those components; 2) scaling the data between 0 and 1 to ensure that decoding was not influenced by the absolute magnitude of the responses. In all cases, a time index of 0 milliseconds indicated the onset of a facial gesture. All facial gestures on a given day were included, except those which were not preceded by at least 1 second of stillness (in order to allow for examination of the premovement period). Decoder performance (accuracy) was estimated by stratified 8-fold cross validation. The folds always referred to the same trials in each window throughout the trial. To determine chance decoding accuracy, we generated a null distribution of 100 samples by permuting the categorical facial gesture labels on all trials and recalculating the 8-fold cross validated decoding accuracy. Significance threshold determined by the 98th percentile of these one hundred permuted decoding accuracies. Note this chance level is not 33% as the three gestures occurred at unequal frequencies; we chose to use all data, permute labels and repeat decoding, rather than sub-select trials. This process was applied independently to each of the pseudopopulations; in **Figure 4B**, mean decoding accuracy (and associated error) in a given time bin was calculated by averaging the decoding performance of 50 pseudopopulations per cortical region in that time bin. Using the full population of neurons resulted in qualitatively and statistically similar results.

### Cross-Temporal Decoding

To assess the stability of neural coding, we applied the above decoding approach to cross-temporal generalization over distinct time points. Again, input to the classifier at each time point was an nTrials × nCells matrix of spike counts, binned at 400 millisecond time windows with 10 millisecond overlap. A smaller overlap window was chosen so as not to potentially obscure fast changes in coding stability. Light denoising and preprocessing was performed similarly as in the categorical decoding analysis. A linear SVM decoder (Matlab *fitcecoc*) was trained at each time window *t* of the trial, and then tested against all other time windows, *t’*, across the trial. By systematically training and testing on all timepoints, we generated a temporal generalization matrix for each cortical region, in which each row corresponds to the time window at which the classifier was trained, and the columns, to the time windows tested. Importantly, the diagonal of this temporal generalization matrix reflects the same performance as the decoder trained and tested only on equivalent time points. Decoder performance was estimated by stratified 8-fold cross validation. Chance decoding accuracy was determined as above. This process was repeated for each of the fifty pseudopopulations in each cortical region. in **Figure 5**, mean decoding accuracy (a single entry in the full matrix) was calculated by averaging the decoding performance of 50 pseudopopulations per cortical region at that *t/t’* timebin. Using the full population of neurons resulted in qualitatively and statistically similar results. Above-chance cross-temporal decoding provides evidence for time-stable population coding. Conversely, failure to generalize across time provides evidence for temporally-dynamic population coding.

### Neural Pairwise Correlations, Population Structure Across Facial Expressions

To assess the structure of neural population activity across cortical regions and facial expressions, we computed a dissimilarity matrix for each (n = 4 cortical regions x 3 gestures = 12). For each facial expression, we first arranged neural activity into a matrix *F* ∈ ℝ ^N x T^, where N is the number of neurons, and T is the number of timepoints. Each cell’s activity during a given facial gesture was represented as its mean-centered average firing rate across all repetitions of that gesture from −500 to +1000 ms (binned at 20 ms, smoothed with 50 ms Gaussian kernel). Dissimilarity was quantified by the correlation distance, *1 – R* (Pearson’s correlation), where correlation is computed between rows of *F* (eg., cells’ activity over time). Each value in the dissimilarity matrix represents a pairwise comparison of two cells’ activity patterns during one type of facial gesture, and indicates the degree to which the trial-averaged response pattern was dissimilar for the two neurons during that facial gesture. When using a common ordering among all matrices (**Figure 3D**, bottom row), ordering of cells/rows was determined via hierarchical clustering of the lipsmacking matrix and applied to all others. Correlation values for all neurons pairs were highly dispersed, indicating the relationship between a pair of neurons during one gesture did not predict their relationship during another. **Figure 3F** depicts per cortical region, the distribution of average R^2^ values (averaged across the three gesture comparisons; threat v. lipsmack, lipsmack v. chew, threat v. chew).

### Trial-Averaged Neural Trajectories

We arranged neural activity into a matrix *M* ∈ ℝ ^FT x N^, where N is the number of neurons, T is the number of timepoints, and *F* is the type of facial gesture. Each cell’s activity during a given facial gesture was represented as its average firing rate across all repetitions of that gesture in the 1000 milliseconds prior to, and after onset of movement (binned at 20 ms, smoothed with 50 ms Gaussian kernel). We performed PCA using singular value decomposition on the mean-centered and standardized *M*, handling each neuron as a variable, to obtain the *N x N* matrix of face gesture PCs. Thus the neural activity at each time point *T*, and for each expression *F*, is represented as a linear combination of face-gesture PCs, where each PC defines a direction in the full *N*-dimensional neural space.

A trajectory’s velocity is a metric which captures the rate of change in neural activation state during that gesture. To determine the velocity of facial gesture trajectories in neural state space, we then calculated the multidimensional Euclidian distance between two successive time bins in N-dimensional neural space for each gesture specific trajectory (**Figure 5** depicts twelve dimensions, similar qualitative results obtained for 4D, 8D, and 20D). Velocities calculated for each of the 3 trajectories were also averaged to provide a single estimation for each cortical region (**Figure 5**, right).

### Decoding of Continuous Facial Kinematics, using Preferential Subspace Identification

To quantify the relationship between neural activity and facial movement, we used preferential subspace identification (PSID) to extract and model behaviorally-relevant neural dynamics in order to predict continuous facial kinematics from high-dimensional neural activity. Once trained, we used the model to perform continuous kinematic decoding of the facial markers and compare performance across cortical regions. The motivation for adopting this method was two-fold: 1) in addition to encoding facial gestures, dynamics in recorded neural activity may also reasonably encode other brain functions (eg., visual responsivity in premotor regions), inputs from other neurons/brain regions (e.g., reciprocal connectivity in M1-S1), and internal states with brain-wide representations (e.g., hunger or affect). These various additional dynamics (which do not directly encode the measured behavior, thus “behaviorally-irrelevant”) are of secondary interest, and may mask or confound behaviorally-relevant dynamics, making their separation key. Secondly, 2) facial gestures are a natural behavior with inherent temporal structure, making a model which explicitly characterizes dynamics, superior to non-dynamic alternatives.

PSID provides such a framework: the state of the brain at each point in time is conceptualized as a high-dimensional latent variable of which some dimensions may drive the measured behavior, some may drive the recorded neural activity, and some may drive both. To directly learn the shared dynamics, the algorithm uses neural activity and behavior jointly during training to identify a general dynamic linear state space model, while prioritizing dynamics relevant to a behavior of interest (here, facial movement). This is achieved by extracting latent states directly from the data, via an orthogonal projection of future behavior onto past neural activity. Once the latent states are extracted, all remaining parameters of the dynamical model are identified via least squares. After the model is trained, an associated Kalman filter operates on novel neural data alone, to generate behavioral and latent state predictions. One important feature of PSID that distinguishes it from related targeted-dimensionality reduction techniques (eg., demixed PCA, others) is its applicability to continuous rather than categorical behavior. Formally, the dynamic linear state space model is formulated as:

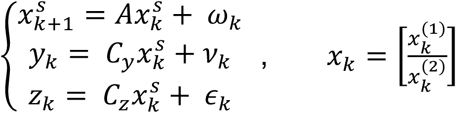

Where:

*k* is time index

9_k_ ∈ ℝ *^n^_y_* is recorded neural activity

*y_k_* ∈ ℝ *^n^_z_* is recorded behavior

*z_k_* ∈ ℝ *^n^_x_* is the latent state driving recorded neural activity, composed of both its behaviorally-relevant (*x_k_* ^(1)^ ∈ ℝ^n^_1_) and irrelevant components (*x_k_* ^(2)^ ∈ ℝ ^n^_2_, with *n_2_* = *n_x_* - *n_1_*).

*A, C_y_, C_z_* are the parameters of the dynamical model learned in training

*ε*_k_ represents behavioral variation in *z_k_*, not represented in the neural data

*w_k_* and *v_k_* are zero-mean white noises, independent of *x_k_*.

PSID has a novel two-stage identification approach that allows it to learn 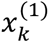 (“behaviorally-relevant” dynamics) directly from training data in its first stage without the need to also learn 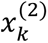 (“behaviorally-irrelevant” dynamics) until an optional second stage. Hyperparameters include *n_1_*, the number of behaviorally-relevant states to be extracted in the first phase, *n_x_*, or the total number of latent states, and *i,* or the horizon, a parameter which affects intermediate linear algebra operations during learning but not final model structure.

The model was applied to our data in the following manner: for a subset of the recordings (2 recording days of subject 1, and 2 recording days of subject 2, totaling 293 unique trials), all simultaneously recorded cells’ firing rates and facial marker data were jointly examined. Data were binned at 50 milliseconds, from −500 milliseconds to +1500 milliseconds around gesture-onset, and smoothed with a 50 millisecond kernel. As in XXX analysis above, we organized the data into pseudopopulations to facilitate comparison of normalized decoding performance across regions. One pseudopopulation was composed of fifty cells from a given cortical region, drawn without replacement from the overall pool of all available cells. The neural activity time series *y_k_* was represented by an array of nTrials, each of which contained a matrix of firing rates of dimensions nTimepoints x nCells. For each cell chosen, all available trials were used. The behavioral time series *z_k_* was represented analogously; rather than using *X* and *Y* positions for each facial marker as the behavior of interest, we performed PCA to obtain the top N principle facial gesture movement components (‘gesture components’) as in FIGURE 1X (see xxx above). Each pseudopopulation therefore contained a unique combination of neurons and their associated behavioral trials. We created twenty-five pseudopopulations, and performed model fitting on each independently. Average decoding performance, defined as the cross-correlation between actual and predicted gesture components on individual trials, was then estimated by stratified, nested k-folds validation procedure (4 outer folds, 8 inner folds), for each model independently in the following manner:

Optimal hyperparameters (*n_1_*, the number of behaviorally-relevant states to be extracted in the first phase; *n_x_*, the total number of latent states; *i,* horizon) were identified via a grid search in the inner loop. The optimal number of behaviorally-relevant states, *n_1_*, was identified by plotting kinematic decoding performance versus *n_x_* when *n_1_* = *n_x_* (i.e., only the first stage of the model fitted) and choosing the smallest *n_1_* which reached within one SEM of peak kinematic decoding performance. The optimal number of total latent states, *n_x_*, was then identified by plotting neural self-prediction performance versus *n_x_* values (for models learned with the optimal *n_1_*) and choosing the smallest *n_x_* which reached within one SEM of peak neural self-prediction performance. This approach favors a lower-dimensional model over a higher-dimensional one with minimal performance gains. In the last two steps, we retrained and evaluated 1) a final two-stage model with the optimal *n_1_*, *n_x_,* and *i,* and 2) a final one-stage model (where *n_1_* = *n_x_*), all within the outer loop. The decoding performances depicted in Figure 3A represent the average performance of all twenty-five pseudopopulations per cortical region; error bars likewise represent +/- 2 standard deviations across iterations.

Importantly, this approach does not force low dimensional dynamics in all cortical regions, but still allows for direct comparison of decoding performances. Finally, data and thus model estimates are based on single-trials; all trial-average estimates were performed ad-hoc as summary metrics for illustrative purposes.

To establish chance levels of decoding performance, each region was compared to a null dataset in which behavioral time series were permuted, and the entire procedure repeated for twenty five permutations. This creates a null dataset which preserves a number of key characteristics of the original while obliterating the relationship between latent neural dynamics and behavior.

## References Cited

1. Parr, L. A., Waller, B. M. & Fugate, J. Emotional communication in primates: implications for neurobiology. Curr. Opin. Neurobiol. 15, 716–720 (2005).

2. Anderson, D. J. & Adolphs, R. A Framework for Studying Emotions across Species. Cell 157, 187–200 (2014).

3. Kret, M. E., Prochazkova, E., Sterck, E. H. M. & Clay, Z. Emotional expressions in human and non-human great apes. Neurosci. Biobehav. Rev. 115, 378–395 (2020).

4. Freiwald, W. A. The Neural Mechanisms of Face Processing: Cells, Areas, Networks, and Models. Curr. Opin. Neurobiol. 60, 184–191 (2020).

5. Yang, Z. & Freiwald, W. A. Encoding of dynamic facial information in the middle dorsal face area. Proc. Natl. Acad. Sci. 120, e2212735120 (2023).

6. Yang, Z. & Freiwald, W. A. Joint encoding of facial identity, orientation, gaze, and expression in the middle dorsal face area. Proc. Natl. Acad. Sci. 118, e2108283118 (2021).

7. Pitcher, D. & Ungerleider, L. G. Evidence for a third visual pathway specialized for social perception. Trends Cogn. Sci. 25, 100–110 (2021).

8. Jenny, A. B. & Saper, C. B. Organization of the facial nucleus and corticofacial projection in the monkey: A reconsideration of the upper motor neuron facial palsy. Neurology 37, 930– 930 (1987).

9. Morecraft, R. J., Louie, J. L., Herrick, J. L. & Stilwell-Morecraft, K. S. Cortical innervation of the facial nucleus in the non-human primate. Brain 124, 176–208 (2001).

10. Morecraft, R. J., Stilwell–Morecraft, K. S. & Rossing, W. R. The Motor Cortex and Facial Expression:: New Insights From Neuroscience. The Neurologist 10, 235–249 (2004).

11. Sobinov, A. R. & Bensmaia, S. J. The neural mechanisms of manual dexterity. Nat. Rev. Neurosci. 22, 741–757 (2021).

12. Burrows, A. M., Waller, B. M. & Parr, L. A. Facial musculature in the rhesus macaque (Macaca mulatta): evolutionary and functional contexts with comparisons to chimpanzees and humans. J. Anat. 215, 320–334 (2009).

13. Waller, B. M., Parr, L. A., Gothard, K. M., Burrows, A. M. & Fuglevand, A. J. Mapping the contribution of single muscles to facial movements in the rhesus macaque. Physiol. Behav. 95, 93–100 (2008).

14. Kavanagh, E., Kimock, C., Whitehouse, J., Micheletta, J. & Waller, B. M. Revisiting Darwin’s comparisons between human and non-human primate facial signals. Evol. Hum. Sci. 4, (2022).

15. MacAllister, R. P., Heagerty, A. & Coleman, K. Behavioral predictors of pairing success in rhesus macaques (Macaca mulatta). Am. J. Primatol. 82, e23081 (2020).

16. Maestripieri, D. Gestural communication in three species of macaques (*Macaca mulatta*, *M. nemestrina*, *M. arctoides*): Use of signals in relation to dominance and social context. Gesture 5, 57–73 (2005).

17. Waller, B. M., Whitehouse, J. & Micheletta, J. Macaques can predict social outcomes from facial expressions. Anim. Cogn. 19, 1031–1036 (2016).

18. Maestripieri, D. & Wallen, K. Affiliative and submissive communication in rhesus macaques. Primates 38, 127–138 (1997).

19. Micheletta, J., Engelhardt, A., Matthews, L., Agil, M. & Waller, B. M. Multicomponent and Multimodal Lipsmacking in Crested Macaques (Macaca nigra). Am. J. Primatol. 75, 763– 773 (2013).

20. Hinde, R. A. & Rowell, T. E. COMMUNICATION BY POSTURES AND FACIAL EXPRESSIONS IN THE RHESUS MONKEY (*MACACA MULATTA*). Proc. Zool. Soc. Lond. 138, 1–21 (1962).

21. Petersen, R. M., Dubuc, C. & Higham, J. P. Facial Displays of Dominance in Non-human Primates. in The Facial Displays of Leaders (ed. Senior, C.) 123–143 (Springer International Publishing, Cham, 2018). doi:10.1007/978-3-319-94535-4_6.

22. Gothard, K. M., Erickson, C. A. & Amaral, D. G. How do rhesus monkeys (Macaca mulatta) scan faces in a visual paired comparison task? Anim. Cogn. 7, 25–36 (2004).

23. Shepherd, S. V., Lanzilotto, M. & Ghazanfar, A. A. Facial Muscle Coordination in Monkeys During Rhythmic Facial Expressions and Ingestive Movements. J. Neurosci. 32, 6105–6116 (2012).

24. Mathis, A. et al. DeepLabCut: markerless pose estimation of user-defined body parts with deep learning. Nat. Neurosci. 21, 1281–1289 (2018).

25. Smith, J. C., Abdala, A. P. L., Borgmann, A., Rybak, I. A. & Paton, J. F. R. Brainstem respiratory networks: building blocks and microcircuits. Trends Neurosci. 36, 152–162 (2013).

26. Morquette, P. et al. Generation of the masticatory central pattern and its modulation by sensory feedback. Prog. Neurobiol. 96, 340–355 (2012).

27. Müri, R. M. Cortical control of facial expression: Cortical Control of Facial Expression. J. Comp. Neurol. 524, 1578–1585 (2016).

28. Arce-McShane, F. I., Hatsopoulos, N. G., Lee, J.-C., Ross, C. F. & Sessle, B. J. Modulation Dynamics in the Orofacial Sensorimotor Cortex during Motor Skill Acquisition. J. Neurosci. 34, 5985–5997 (2014).

29. Arce-McShane, F. I., Sessle, B. J., Ram, Y., Ross, C. F. & Hatsopoulos, N. G. Multiple regions of sensorimotor cortex encode bite force and gape. Front. Syst. Neurosci. 17, 1213279 (2023).

30. Lin, L. D., Murray, G. M. & Sessle, B. J. Functional properties of single neurons in the primate face primary somatosensory cortex. I. Relations with trained orofacial motor behaviors. J. Neurophysiol. 71, 2377–2390 (1994).

31. Sessle, B. J. et al. Properties and plasticity of the primate somatosensory and motor cortex related to orofacial sensorimotor function. Clin. Exp. Pharmacol. Physiol. 32, 109–114 (2005).

32. Bouras, T., Stranjalis, G. & Sakas, D. E. Traumatic midbrain hematoma in a patient presenting with an isolated palsy of voluntary facial movements: Case report. J. Neurosurg. 107, 158–160 (2007).

33. Cerrato, P. et al. Emotional facial paresis in a patient with a lateral medullary infarction. Neurology 60, 723–724 (2003).

34. Dawson, K., Hourihan, M. D., Wiles, C. M. & Chawla, J. C. Separation of voluntary and limbic activation of facial and respiratory muscles in ventral pontine infarction. J. Neurol. Neurosurg. Psychiatry 57, 1281–1282 (1994).

35. Holstege, G. Emotional innervation of facial musculature. Mov. Disord. 17, S12–S16 (2002).

36. Hopf, H. C., Fitzek, C., Marx, J., Urban, P. P. & Thömke, F. Emotional facial paresis of pontine origin. Neurology 54, 1217 (2000).

37. Michel, L. et al. Emotional facial palsy following striato-capsular infarction. J. Neurol. Neurosurg. Psychiatry 79, 193–4 (2008).

38. Topper, R., Kosinski, C. & Mull, M. Volitional type of facial palsy associated with pontine ischaemia. J. Neurol. Neurosurg. Psychiatry 58, 732–734 (1995).

39. Trepel, M., Weller, M., Dichgans, J. & Petersen, D. Voluntary facial palsy with a pontine lesion. J. Neurol. Neurosurg. Psychiatry 61, 531–533 (1996).

40. Raos, V., Umiltá, M.-A., Murata, A., Fogassi, L. & Gallese, V. Functional Properties of Grasping-Related Neurons in the Ventral Premotor Area F5 of the Macaque Monkey. J. Neurophysiol. 95, 709–729 (2006).

41. Sani, O. G., Abbaspourazad, H., Wong, Y. T., Pesaran, B. & Shanechi, M. M. Modeling behaviorally relevant neural dynamics enabled by preferential subspace identification. Nat. Neurosci. 24, 140–149 (2021).

42. Gallego, J. A. et al. Cortical population activity within a preserved neural manifold underlies multiple motor behaviors. Nat. Commun. 9, 4233 (2018).

43. Gallego, J. A., Perich, M. G., Miller, L. E. & Solla, S. A. Neural Manifolds for the Control of Movement. Neuron 94, 978–984 (2017).

44. Parr, L. A., Waller, B. M., Burrows, A. M., Gothard, K. M. & Vick, S. J. Brief communication: MaqFACS: A muscle-based facial movement coding system for the rhesus macaque. Am. J. Phys. Anthropol. 143, 625–630 (2010).

45. Partan, S. R. Single and Multichannel Signal Composition: Facial Expressions and Vocalizations of Rhesus Macaques (Macaca mulatta). Behaviour 139, 993–1027 (2002).

46. Cunningham, J. P. & Yu, B. M. Dimensionality reduction for large-scale neural recordings. Nat. Neurosci. 17, 1500–1509 (2014).

47. King, J.-R. & Dehaene, S. Characterizing the dynamics of mental representations: the temporal generalization method. Trends Cogn. Sci. 18, 203–210 (2014).

48. Meyers, E. M., Freedman, D. J., Kreiman, G., Miller, E. K. & Poggio, T. Dynamic Population Coding of Category Information in Inferior Temporal and Prefrontal Cortex. J. Neurophysiol. 100, 1407–1419 (2008).

49. Ceccarelli, F. et al. Static and dynamic coding in distinct cell types during associative learning in the prefrontal cortex. Nat. Commun. 14, 8325 (2023).

50. Enel, P., Wallis, J. D. & Rich, E. L. Stable and dynamic representations of value in the prefrontal cortex. eLife 9, e54313 (2020).

51. Oh, B.-I., Kim, Y.-J. & Kang, M.-S. Ensemble representations reveal distinct neural coding of visual working memory. Nat. Commun. 10, 5665 (2019).

52. Sapountzis, P., Paneri, S., Papadopoulos, S. & Gregoriou, G. G. Dynamic and stable population coding of attentional instructions coexist in the prefrontal cortex. Proc. Natl. Acad. Sci. 119, e2202564119 (2022).

53. Stokes, M. G. et al. Dynamic Coding for Cognitive Control in Prefrontal Cortex. Neuron 78, 364–375 (2013).

54. Tsao, D. Y. A Cortical Region Consisting Entirely of Face-Selective Cells. Science 311, 670– 674 (2006).

55. Landi, S. M. & Freiwald, W. A. Two areas for familiar face recognition in the primate brain. Science 357, 591–595 (2017).

56. Charlesworth & Kreutzer, W. R. & M. A. Darwin and Facial Expression: A Century of Research in Review. in Facial expressions of infants and children 91–168 (1973).

57. Ekman, P. Facial expression and emotion. Am. Psychol. 48, 384–392 (1993).

58. Pehlevan, C., Ali, F. & Ölveczky, B. P. Flexibility in motor timing constrains the topology and dynamics of pattern generator circuits. Nat. Commun. 9, 977 (2018).

59. Hopf, H. C., Md, W. M.-F. & Hopf, N. J. Localization of emotional and volitional facial paresis. Neurology 42, 1918–1918 (1992).

60. Kappos, L. & Mehling, M. Dissociation of Voluntary and Emotional Innervation after Stroke. N. Engl. J. Med. 363, e25 (2010).

61. Ross, R. T. & Mathiesen, R. Volitional and Emotional Supranuclear Facial Weakness. N. Engl. J. Med. 338, 1515–1515 (1998).

62. Carr, L., Iacoboni, M., Dubeau, M.-C., Mazziotta, J. C. & Lenzi, G. L. Neural mechanisms of empathy in humans: A relay from neural systems for imitation to limbic areas. Proc. Natl. Acad. Sci. 100, 5497–5502 (2003).

63. Hennenlotter, A. et al. A common neural basis for receptive and expressive communication of pleasant facial affect. NeuroImage 26, 581–591 (2005).

64. Iwase, M. et al. Neural Substrates of Human Facial Expression of Pleasant Emotion Induced by Comic Films: A PET Study. NeuroImage 17, 758–768 (2002).

65. Krippl, M., Karim, A. A. & Brechmann, A. Neuronal correlates of voluntary facial movements. Front. Hum. Neurosci. 9, (2015).

66. Leslie, K. R., Johnson-Frey, S. H. & Grafton, S. T. Functional imaging of face and hand imitation: towards a motor theory of empathy. NeuroImage 21, 601–607 (2004).

67. Wild, B. et al. Humor and smiling: Cortical regions selective for cognitive, affective, and volitional components. Neurology 66, 887–893 (2006).

68. Shepherd, S. V. & Freiwald, W. A. Functional Networks for Social Communication in the Macaque Monkey. Neuron 99, 413–420.e3 (2018).

69. Churchland, M. M. & Shenoy, K. V. Temporal complexity and heterogeneity of single-neuron activity in premotor and motor cortex. J. Neurophysiol. 97, 4235–4257 (2007).

70. Shenoy, K. V., Sahani, M. & Churchland, M. M. Cortical Control of Arm Movements: A Dynamical Systems Perspective. Annu. Rev. Neurosci. 36, 337–359 (2013).

71. Wang, T., Chen, Y. & Cui, H. From Parametric Representation to Dynamical System: Shifting Views of the Motor Cortex in Motor Control. Neurosci. Bull. 38, 796–808 (2022).

72. Cattaneo, L., Saccani, E., De Giampaulis, P., Crisi, G. & Pavesi, G. Central facial palsy revisited: A clinical-radiological study. Ann. Neurol. 68, 404–408 (2010).

73. Arnts, H. et al. On the pathophysiology and treatment of akinetic mutism. Neurosci. Biobehav. Rev. 112, 270–278 (2020).

74. N□meth, G., Heged□s, K. & Moln□r, L. Akinetic mutism associated with bicingular lesions: Clinicopathological and functional anatomical correlates. Eur. Arch. Psychiatry Neurol. Sci. 237, 218–222 (1988).

75. Kim, C. M., Finkelstein, A., Chow, C. C., Svoboda, K. & Darshan, R. Distributing task-related neural activity across a cortical network through task-independent connections. Nat. Commun. 14, 2851 (2023).

76. Elsayed, G. F., Lara, A. H., Kaufman, M. T., Churchland, M. M. & Cunningham, J. P. Reorganization between preparatory and movement population responses in motor cortex. Nat. Commun. 7, 13239 (2016).

77. Kaufman, M. T., Churchland, M. M., Ryu, S. I. & Shenoy, K. V. Cortical activity in the null space: permitting preparation without movement. Nat. Neurosci. 17, 440–448 (2014).

78. Perich, M. G. et al. Motor cortical dynamics are shaped by multiple distinct subspaces during naturalistic behavior. 2020.07.30.228767 Preprint at 10.1101/2020.07.30.228767 (2020).

79. Stavisky, S. D., Kao, J. C., Ryu, S. I. & Shenoy, K. V. Motor Cortical Visuomotor Feedback Activity Is Initially Isolated from Downstream Targets in Output-Null Neural State Space Dimensions. Neuron 95, 195–208.e9 (2017).

80. Dekleva, B. M. et al. Motor cortex retains and reorients neural dynamics during motor imagery. Nat. Hum. Behav. 1–14 (2024) doi:10.1038/s41562-023-01804-5.

81. Haggard, M. & Chacron, M. J. Nonresponsive Neurons Improve Population Coding of Object Location. J. Neurosci. 45, e1068242024 (2025).

82. Bouchard, K. E. & Chang, E. F. Control of Spoken Vowel Acoustics and the Influence of Phonetic Context in Human Speech Sensorimotor Cortex. J. Neurosci. 34, 12662–12677 (2014).

83. Chartier, J., Anumanchipalli, G. K., Johnson, K. & Chang, E. F. Encoding of Articulatory Kinematic Trajectories in Human Speech Sensorimotor Cortex. Neuron 98, 1042–1054.e4 (2018).

84. Mazurek, K. A., Rouse, A. G. & Schieber, M. H. Mirror Neuron Populations Represent Sequences of Behavioral Epochs During Both Execution and Observation. J. Neurosci. 38, 4441–4455 (2018).

85. Lara, A. H., Cunningham, J. P. & Churchland, M. M. Different population dynamics in the supplementary motor area and motor cortex during reaching. Nat. Commun. 9, (2018).

86. Aflalo, T. N. & Graziano, M. S. A. Partial tuning of motor cortex neurons to final posture in a free-moving paradigm. Proc. Natl. Acad. Sci. 103, 2909–2914 (2006).

87. Shenoy, K. V. & Carmena, J. M. Combining Decoder Design and Neural Adaptation in Brain-Machine Interfaces. Neuron 84, 665–680 (2014).

88. Aggarwal, V., Mollazadeh, M., Davidson, A. G., Schieber, M. H. & Thakor, N. V. State-based decoding of hand and finger kinematics using neuronal ensemble and LFP activity during dexterous reach-to-grasp movements. J. Neurophysiol. 109, 3067–3081 (2013).

89. Tanji, J. & Evarts, E. V. Anticipatory activity of motor cortex neurons in relation to direction of an intended movement. J. Neurophysiol. 39, 1062–1068 (1976).

90. Cisek, P. Integrated Neural Processes for Defining Potential Actions and Deciding between Them: A Computational Model. J. Neurosci. 26, 9761–9770 (2006).

91. Bastian, A., Schöner, G. & Riehle, A. Preshaping and continuous evolution of motor cortical representations during movement preparation. Eur. J. Neurosci. 18, 2047–2058 (2003).

92. Afshar, A. et al. Single-Trial Neural Correlates of Arm Movement Preparation. Neuron 71, 555–564 (2011).

93. Ames, K. C., Ryu, S. I. & Shenoy, K. V. Neural Dynamics of Reaching following Incorrect or Absent Motor Preparation. Neuron 81, 438–451 (2014).

94. Churchland, M. M., Cunningham, J. P., Kaufman, M. T., Ryu, S. I. & Shenoy, K. V. Cortical Preparatory Activity: Representation of Movement or First Cog in a Dynamical Machine? Neuron 68, 387–400 (2010).

95. Sussillo, D., Churchland, M. M., Kaufman, M. T. & Shenoy, K. V. A neural network that finds a naturalistic solution for the production of muscle activity. Nat. Neurosci. 18, 1025– 1033 (2015).

96. Zinger, N., Harel, R., Gabler, S., Israel, Z. & Prut, Y. Functional Organization of Information Flow in the Corticospinal Pathway. J. Neurosci. 33, 1190–1197 (2013).

97. Harel, R. et al. Computation in spinal circuitry: Lessons from behaving primates. Behav. Brain Res. 194, 119–128 (2008).

98. Dum, R. P. & Strick, P. L. Motor areas in the frontal lobe of the primate. Physiol. Behav. 77, 677–682 (2002).

99. Dum, R. & Strick, P. The origin of corticospinal projections from the premotor areas in the frontal lobe. J. Neurosci. 11, 667–689 (1991).

100. Kurtzer, I., Herter, T. M. & Scott, S. H. Random change in cortical load representation suggests distinct control of posture and movement. Nat. Neurosci. 8, 498–504 (2005).

101. Stark, E., Asher, I. & Abeles, M. Encoding of Reach and Grasp by Single Neurons in Premotor Cortex Is Independent of Recording Site. J. Neurophysiol. (2007) doi:10.1152/jn.01328.2006.

102. Flash, T. & Hogan, N. The coordination of arm movements: an experimentally confirmed mathematical model. J. Neurosci. 5, 1688–1703 (1985).

103. Merel, J., Botvinick, M. & Wayne, G. Hierarchical motor control in mammals and machines. Nat. Commun. 10, 5489 (2019).

104. Parthasarathy, A. et al. Mixed selectivity morphs population codes in prefrontal cortex. Nat. Neurosci. 20, 1770–1779 (2017).

105. Snyder, A. C., Yu, B. M. & Smith, M. A. A Stable Population Code for Attention in Prefrontal Cortex Leads a Dynamic Attention Code in Visual Cortex. J. Neurosci. 41, 9163–9176 (2021).

106. Murray, J. D. et al. A hierarchy of intrinsic timescales across primate cortex. Nat. Neurosci. 17, 1661–1663 (2014).

107. Cavanagh, S. E., Towers, J. P., Wallis, J. D., Hunt, L. T. & Kennerley, S. W. Reconciling persistent and dynamic hypotheses of working memory coding in prefrontal cortex. Nat. Commun. 9, 3498 (2018).

108. Chaudhuri, R., Knoblauch, K., Gariel, M.-A., Kennedy, H. & Wang, X.-J. A Large-Scale Circuit Mechanism for Hierarchical Dynamical Processing in the Primate Cortex. Neuron 88, 419–431 (2015).

109. Spitmaan, M., Seo, H., Lee, D. & Soltani, A. Multiple timescales of neural dynamics and integration of task-relevant signals across cortex. Proc. Natl. Acad. Sci. 117, 22522–22531 (2020).

110. Metzger, S. L. et al. A high-performance neuroprosthesis for speech decoding and avatar control. Nature 620, 1037–1046 (2023).

111. Willett, F. R. et al. A high-performance speech neuroprosthesis. Nature 620, 1031–1036 (2023).

112. Card, N. S. et al. An Accurate and Rapidly Calibrating Speech Neuroprosthesis. N. Engl. J. Med. 391, 609–618 (2024).

113. Dum, R. P. & Strick, P. L. The origin of corticospinal projections from the premotor areas in the frontal lobe. J. Neurosci. 11, 667–689 (1991).

114. Picard, N. & Strick, P. L. Motor Areas of the Medial Wall: A Review of Their Location and Functional Activation. Cereb. Cortex 6, 342–353 (1996).

115. Morecraft, R. J. & Van Hoesen, G. W. Convergence of Limbic Input to the Cingulate Motor Cortex in the Rhesus Monkey. Brain Res. Bull. 45, 209–232 (1998).

116. Bates, J. F. & Goldman-Rakic, P. S. Prefrontal connections of medial motor areas in the rhesus monkey. J. Comp. Neurol. 336, 211–228 (1993).

117. Morecraft, R. J. et al. Amygdala interconnections with the cingulate motor cortex in the rhesus monkey. J. Comp. Neurol. 500, 134–165 (2007).

118. Shima, K. et al. Two movement-related foci in the primate cingulate cortex observed in signal-triggered and self-paced forelimb movements. J. Neurophysiol. 65, 188–202 (1991).

119. Crutcher, M. D., Russo, G. S., Ye, S. & Backus, D. A. Target-, limb-, and context-dependent neural activity in the cingulate and supplementary motor areas of the monkey. Exp. Brain Res. 158, 278–288 (2004).

120. Iwata, J., Shima, K., Tanji, J. & Mushiake, H. Neurons in the cingulate motor area signal context-based and outcome-based volitional selection of action. Exp. Brain Res. 229, 407– 417 (2013).

121. Shima, K. & Tanji, J. Role for Cingulate Motor Area Cells in Voluntary Movement Selection Based on Reward. Science 282, 1335–1338 (1998).

122. Williams, Z. M., Bush, G., Rauch, S. L., Cosgrove, G. R. & Eskandar, E. N. Human anterior cingulate neurons and the integration of monetary reward with motor responses. Nat. Neurosci. 7, 1370–1375 (2004).

123. Sliwa, J. & Freiwald, W. A. A dedicated network for social interaction processing in the primate brain. Science 356, 745–749 (2017).

124. Alagapan, S. et al. Cingulate dynamics track depression recovery with deep brain stimulation. Nature 622, 130–138 (2023).

125. Liao, D. A., Zhang, Y. S., Cai, L. X. & Ghazanfar, A. A. Internal states and extrinsic factors both determine monkey vocal production. Proc. Natl. Acad. Sci. 115, 3978–3983 (2018).

126. Remedios, R. et al. Social behaviour shapes hypothalamic neural ensemble representations of conspecific sex. Nature 550, 388–392 (2017).

127. Wood, A., Rychlowska, M., Korb, S. & Niedenthal, P. Fashioning the Face: Sensorimotor Simulation Contributes to Facial Expression Recognition. Trends Cogn. Sci. 20, 227–240 (2016).

128. Chaure, F. J., Rey, H. G. & Quian Quiroga, R. A novel and fully automatic spike-sorting implementation with variable number of features. J. Neurophysiol. 120, 1859–1871 (2018).

129. Livneh, U., Resnik, J., Shohat, Y. & Paz, R. Self-monitoring of social facial expressions in the primate amygdala and cingulate cortex. Proc. Natl. Acad. Sci. 109, 18956–18961 (2012).

